# WitChi: Efficient Detection and Pruning of Compositional Bias in Phylogenomic Alignments Using Empirical Chi-Squared Testing

**DOI:** 10.1101/2025.07.14.663642

**Authors:** Stephan Köstlbacher, Kassiani Panagiotou, Daniel Tamarit, Thijs J. G. Ettema

## Abstract

Convergent evolution, where unrelated taxa independently evolve similar nucleotide or amino acid compositions, can introduce compositional bias into biological sequence data. Such biases distort phylogenetic inference, particularly in deep or unevenly sampled phylogenomic datasets. While composition-aware models can mitigate this issue, their computational demands often preclude their use in large-scale analyses. We present WitChi, a computationally efficient tool for identifying and removing compositionally biased alignment columns using empirical significance testing. WitChi calculates taxon-specific chi-squared (χ²) scores and compares them to null distributions derived from permutations within alignment columns that preserve the phylogenetic structure of the alignment. Sites most responsible for deviation from the expected null are iteratively pruned using one of three scoring algorithms until the bias is no longer statistically detectable. Z-scores and p-values are provided for both taxa and alignments, offering interpretable metrics of the magnitude of compositional bias. Pruning of simulated compositional heterogeneous alignments show that WitChi reliably restores correct topologies under standard, compositionally stationary models. In benchmarks, WitChi outperforms BMGE’s stationary-based trimming while scaling linearly with taxon number. Applied to the archaeal GTDB r220 dataset (5,869 taxa; 10,101 sites), WitChi completes pruning in under one hour on four CPU cores. The resulting phylogeny recovers key clades previously resolved only by in-depth analyses using complex models of sequence evolution. WitChi provides an efficient, scalable solution for detecting and removing compositional bias in phylogenomic datasets comprising thousands to tens of thousands of taxa, enabling more accurate phylogenetic inference across the tree of life.

## Introduction

Compositional heterogeneity in multiple sequence alignments (MSAs) is a well-recognized source of systematic error in phylogenetic inference (Foster and Hickey 1999). When different lineages evolve under distinct nucleotide or amino acid compositions, phylogenetic methods may recover spurious relationships, grouping taxa by sequence composition rather than evolutionary history. This issue is particularly relevant in large phylogenomic datasets, where compositional biases may be widespread and difficult to detect.

Notable examples include the misplacement of AT-biased (Foster et al. 1997), halophilic (Baker et al. 2024; Martijn et al. 2020; Aouad et al. 2019), and thermophilic lineages (Eme et al. 2023; Huang et al. 2025)—many of which belong to Archaea. This domain, in particular, has proven vulnerable to compositional artefacts due to the ecological extremes inhabited by many of its taxa. In each case, biased composition overwhelmed the true phylogenetic signal, often leading to highly supported yet incorrect topologies without laborious specialized analyses.

To address these challenges, some methods model substitution processes using multiple composition matrices across the tree. For example, non-stationary models implemented in software such as p4 (Foster 2004) explicitly account for compositional heterogeneity across branches, but are computationally demanding. Recoding strategies—such as Dayhoff (Dayhoff et al. 1969) or SR4 schemes (Susko and Roger 2007)—aim to reduce compositional noise by collapsing amino acids into categories with similar physicochemical properties or high frequencies of substitutions, thus eliminating within-category compositional biases. However, these approaches can only target some sources of heterogeneity and are not universally successful at mitigating compositional biases (Foster et al. 2023).

A flexible and widely used alternative is chi-squared (χ²) pruning, which removes alignment columns that disproportionately contribute to taxon-specific compositional biases (Esser et al. 2004; Viklund et al. 2012; Williams et al. 2007). This method does not rely on prior tree inference or predefined recoding, but instead uses taxon-specific χ² scores to identify problematic sites (Viklund et al. 2012; Martijn et al. 2015). It is thus flexible and agnostic to the form of heterogeneity, successfully mitigating diverse sources of compositional bias (Viklund et al. 2012; Martijn et al. 2015, 2020; Dharamshi et al. 2020, 2023; Huang et al. 2025; Martijn et al. 2018; Muñoz-Gómez et al. 2022). A related strategy is implemented in BMGE (Criscuolo and Gribaldo 2010), which includes a stationary-based trimming mode that iteratively removes compositionally deviant sites using pairwise comparisons of sequence composition. Like χ² pruning, this approach is model-free and does not require a tree, yet it lacks a statistical testing framework for taxon-specific bias and scales poorly with large numbers of taxa.

However, a more fundamental limitation remains: χ² tests assume that each sequence is an independent sample—a condition violated in MSAs due to shared ancestry. As a result, the canonical parametric χ² distribution, which assumes independent and identically distributed data, does not provide an accurate baseline for interpreting scores from real MSAs (Goldman 1993). While simulations can effectively generate empirical null distributions (Foster 2004), this approach requires a known tree topology and carefully chosen substitution model. These prerequisites introduce practical limitations, making simulations labor-intensive to set up and computationally costly for large phylogenomic datasets.

To overcome these limitations, we developed WitChi, a fast and modular Python tool that uses empirical null distributions derived from column permutations to test for compositionally biased taxa. WitChi iteratively removes alignment columns that drive taxon-specific χ² scores away from the permuted null expectation and recalculates alignment-wide amino acid frequencies to approach an unbiased equilibrium. This permutation-based framework better reflects the statistical structure of real MSAs and eliminates the need for prior tree inference or simulation. WitChi integrates the established χ² pruning method (“squared”) and introduces two additional scoring algorithms. We compare WitChi to the stationary-based trimming mode in BMGE, showing that WitChi achieves slightly superior clade recovery while scaling linearly—rather than quadratically—with the number of taxa, making it suitable for phylogenomic datasets with hundreds to thousands of taxa. We benchmark the tool on simulated datasets with varying degrees of heterogeneity and apply it to the archaeal tree of life—an empirically well-studied dataset where many taxa show compositionally biased sequences—demonstrating both practical scalability and significant gains in phylogenetic accuracy.

## Implementation and Evaluation

### WitChi Workflow Overview

WitChi is a modular Python-based framework designed to detect and prune compositionally biased columns from MSAs. It operates in two main modes: ‘test’ and ‘prune’ (Fig. 1). These can be run independently or as an iterative workflow. Both rely on the calculation of taxon-specific χ² scores and comparison to an empirical null distribution generated by column-wise permutation.

**Figure 1.**
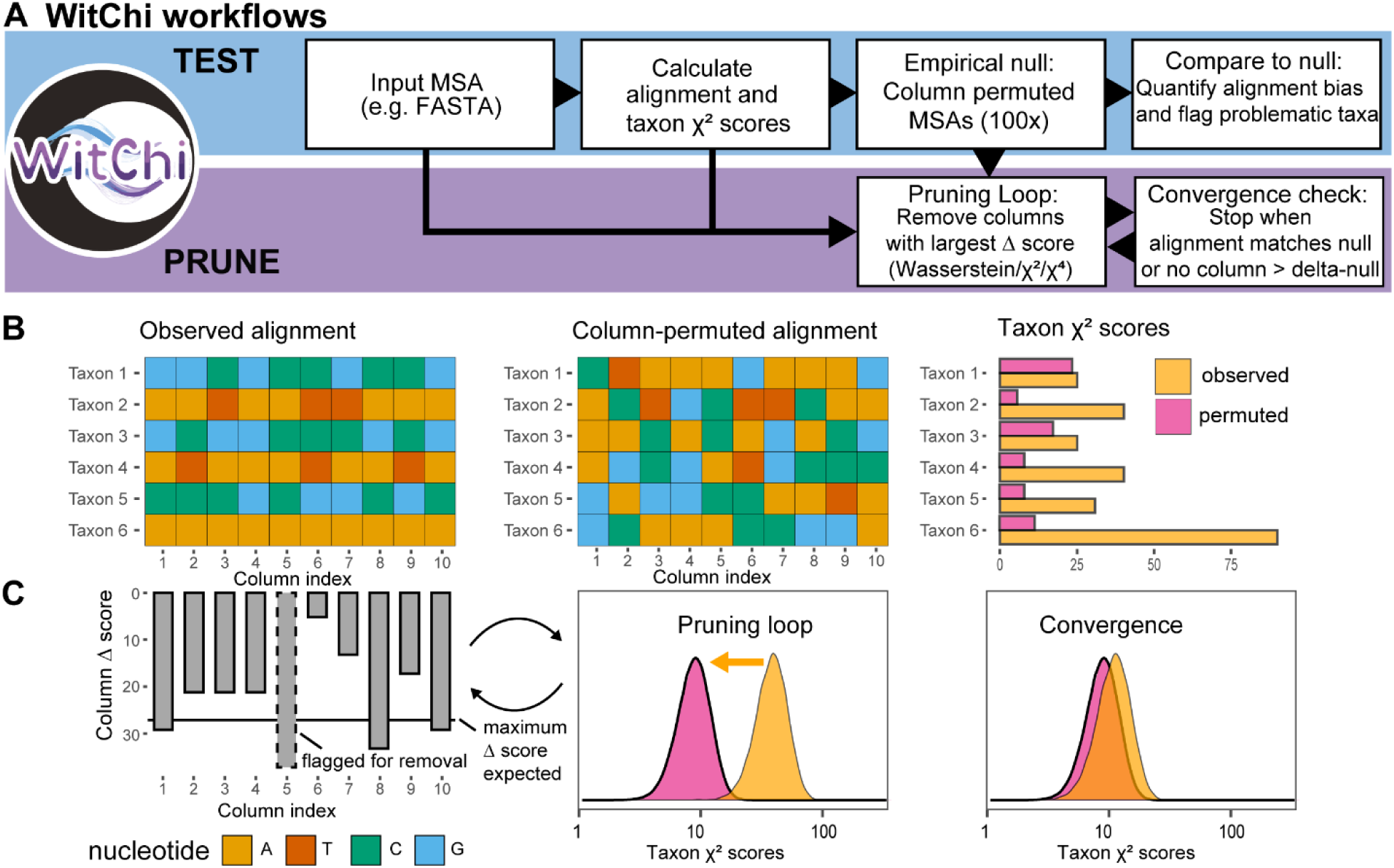
Overview of the WitChi workflows for detecting and reducing compositional bias in multiple sequence alignments. (**A**) The ’test’ workflow (blue background) computes taxon-specific χ² scores and builds an empirical null distribution by column permutation (100×), identifying biased taxa. The ’prune’ workflow (purple background) iteratively removes the alignment columns with the highest Δ-score — defined by the active scoring algorithm (squared, Δχ²; quartic, Δχ⁴; or Wasserstein, ΔW) — followed by a convergence check that halts pruning once the alignment-level empirical p-value exceeds 0.05 or no remaining column’s Δ-score exceeds the per-column delta-null (whichever occurs first). (**B**) An example observed MSA and one corresponding column-permuted MSA, showing how taxon-specific composition is homogenized while the column-wise (global) residue distribution is preserved; the bar plot at right compares taxon χ² scores between the observed and permuted alignments. (**C**) Left: per-column Δ-scores for the observed alignment in (B), with the most biased column flagged for removal (dashed box) when its Δ-score exceeds the maximum expected under the permuted null. Middle (pruning loop): the density of taxon-specific χ² scores shifts from the observed alignment (orange) toward the permuted null (pink) as columns are removed. Right (convergence): once the stopping criterion is met, pruning halts and the observed taxon χ² distribution overlaps the permuted null.

The ‘test’ module computes a χ² score for each taxon, quantifying how much its composition deviates from the alignment-wide average. To account for phylogenetic structure, WitChi generates a null distribution by independently permuting the residues within each column, that is, shuffling character states across taxa separately for each site, so the randomization is independent over sites. This preserves site-specific residue distributions while disrupting taxon-specific signals. The empirical null is used to calculate Z-scores and empirical p-values for each taxon.

The ‘prune’ module removes alignment columns that contribute most disproportionately to bias. At each iteration, columns are scored using one of three algorithms—squared, quartic, or Wasserstein-guided (see below)—and the most extreme columns are iteratively pruned. Unless a user-defined threshold is reached first, pruning continues until the alignment empirical p-value exceeds 0.05 or no column’s Δ-score is distinguishable from the permutation null (see Pruning loop and stopping criteria).

### Software Implementation

WitChi is implemented in Python 3.9 and designed for speed, scalability, extensibility, and ease of use. The core scientific stack includes NumPy for efficient array operations and matrix-based statistics (Harris et al. 2020), Biopython for robust parsing of FASTA, PHYLIP, and NEXUS alignments (Cock et al. 2009), multiprocessing with joblib for parallelized permutation and pruning steps (Varoquaux et al., n.d.). WitChi includes a command-line interface (‘test’ and ‘prune’) with sensible defaults and logging. It is available under the MIT license at https://github.com/stephkoest/witchi, with included unit tests, example data, and benchmark scripts.

### Statistical Basis

To quantify compositional bias in MSAs, we calculate a χ² score for each taxon. This score reflects how much the observed frequency of characters (e.g., nucleotides or amino acids) in a given sequence deviates from the global character distribution of the alignment. Specifically, we compute:

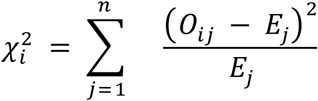

Where *O_i,j_* and *E_j_* are the observed and expected frequencies of amino acid *j* in taxon *i* in the alignment, and *n* is the number of characters (20 amino-acids or 4 nucleotides). Then, we compute an alignment-wide χ² score:

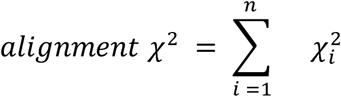

Where *χ_i_²* is the χ² score of taxon *i*, and *n* is the number of taxa in the alignment.

#### Empirical Null Distribution from Permutations

To approximate an appropriate null distribution, we randomly permute characters within each alignment column, maintaining global composition while removing taxon-specific signal. This generates permuted MSAs that reflect the expected χ² score distribution under compositional homogeneity. Each permutation results in a new χ² score for every taxon. Across all permutations (default: 100), these scores are aggregated to form an empirical null distribution. We calculated that P = 100 permutations suffice for the stopping decisions: across 6,000 simulated alignments, the alignment-level and per-column delta-null q95 thresholds at P = 100 differ from their P = 10,000 values only by a median of 2.2% and 2.6% (R² = 0.995 and 0.989), well below the downstream tree-reconstruction variance.

#### Taxon- and Alignment-Level Bias Metrics

To assess compositional bias in multiple sequence alignments, WitChi calculates a suite of taxon-specific and alignment-level statistics based on χ² scores and their empirical null distributions.

##### Taxon-Specific χ² Scores and Empirical Testing

To determine whether a taxon’s observed χ² score indicates significant compositional bias, components are defined as follows with:

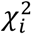 _obs_: the observed taxon-specific χ² score from the input alignment.

χ²_perm_: the set of χ² scores for taxa across N permuted alignments (the empirical null distribution).

μ_perm_ and MAD_perm_: the mean and median absolute deviation of the empirical null distribution over all taxa from the permuted alignments.

##### Taxon empirical p-values

To quantify the significance of each taxon’s observed χ² score, empirical p-values are computed by comparing them to the permuted null and multiplied by N_obs as a Bonferroni correction for the N_obs taxa simultaneously tested for compositional bias, where N_obs is the number of taxa in the alignment.:

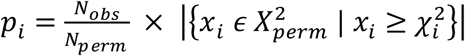

Here, *p_i_* is the empirical p-value for taxon i, *N_perm_* is the number of permuted alignments, and *N_obs_* is the number of observed χ^2^ values. This value reflects how extreme the observed χ² score is under the null hypothesis of compositional homogeneity. We chose an arbitrary cutoff of empirical p-value smaller than 0.05 to suggest significant deviation.

##### Taxon Z-scores

We compute robust Z-scores to express the magnitude of deviation from the empirical null:

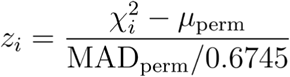

These per-taxon scores can be used to identify outlier taxa in the ‘test’ module of the workflow.

#### Alignment-Level Summary Statistics

##### Z-score

The alignment-level Z is computed analogously from the summed observed χ² across taxa:

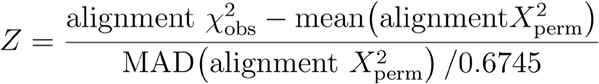

Where alignment χ² = Σᵢ χ²ᵢ and captures how far the alignment scores deviate from expectation.

##### Alignment empirical p-value

To test whether the overall alignment contains significant compositional bias, we compute an empirical p-value for the alignment (summed) χ² score:

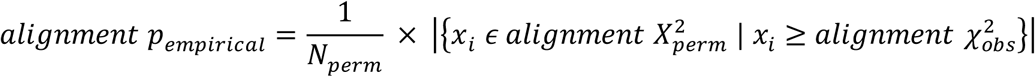

This represents the probability that a randomly permuted alignment would have an equal or greater χ² score than observed.

The minimum Bonferroni-corrected per-taxon p-value is itself a valid omnibus test of homogeneity. We use the alignment-level empirical p-value as our compositional bias criterion because it is robust to tree shape, whereas the per-taxon minimum is built on the per-taxon null, which is sensitive to branch-length heterogeneity (Fig. S1B). The two tests are complementary in power, the per-taxon minimum being most sensitive when bias is concentrated in a few taxa and the alignment-level test when bias is diffuse across many.

### Scoring Algorithms

WitChi implements three scoring strategies to rank alignment columns by their contribution to compositional bias. Each reflects a different principle for identifying columns that distort χ² statistics across taxa.

The Squared scoring algorithm ranks columns by the change in the summed per-taxon χ² (first moment);. The Quartic algorithm ranks columns by the summed square of per-taxon χ² scores, up-weighting across-taxon variance (a column whose deviation is concentrated in few high-deviation taxa scores higher); The Wasserstein algorithm ranks how each column shifts the full distribution of χ² scores relative to the permuted null, capturing subtle or distribution-wide effects.

#### Squared

We compute a total χ^2^ value across all rows. For each column, we determine how its removal affects the global χ^2^ score. Columns producing the largest drop are pruned first.

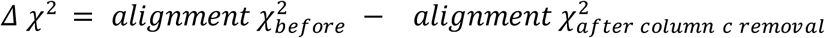

A positive *Δχ*^2^ indicates that pruning has shifted the alignment’s composition closer to the expected distribution under homogeneity.

#### Quartic

We squared the row-level χ^2^ values and then calculated the impact of removing each column on these squared scores. This approach amplifies high values, emphasizing columns that strongly deviate from the expected distribution.

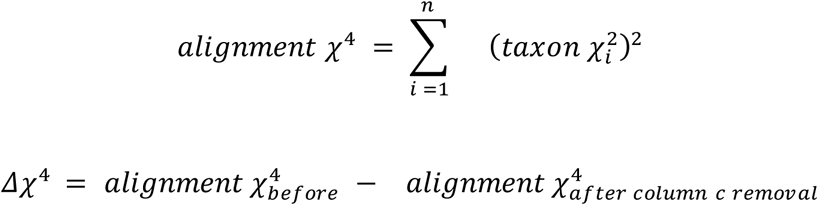

A positive *Δχ*^4^ indicates that pruning has shifted the alignment’s composition closer to the expected distribution under homogeneity.

#### Wasserstein

The Wasserstein-guided scorer ranks columns by how far the observed per-taxon score distribution lies from the homogeneous null. For a column, the *Δ*-score is the average absolute difference between the sorted observed and sorted permuted-null per-taxon Z-scores — that is, the 1-Wasserstein (Kantorovich) distance between the two distributions (Kantorovich 1960). We compute it on matched quantiles of the two distributions, which keeps the scorer fast and comparable across alignments of different size. Let 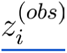 and 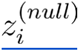 denote the i-th quantiles of the observed and permutation-null per-taxon Z-score distributions over 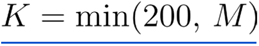 equally spaced positions, where M is the number of null values:

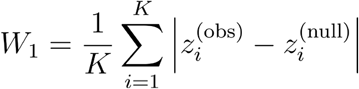

the L1 distance between the two ordered quantile sequences.

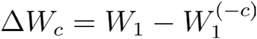

A positive Δ*W_c_* (the decrease in W₁ when a column is removed) indicates that pruning has shifted the observed Z-score distribution closer to the homogeneous null.

### Pruning loop and stopping criteria

In each iteration, the active scoring algorithm ranks all columns by Δ-score and the top k columns (--top_n, default 1) are nominated for removal. Two tests govern whether they are removed.

#### Alignment-level stop

After each batch removal, the alignment empirical p-value (Eq. above) is recomputed on the pruned alignment. Pruning halts if alignment empirical p-value exceeds 0.05.

#### Per-column delta-null test

We generate Pₙᵤₗₗ = --permutations additional permuted alignments (default 100), seeded by 12345 + N_perm + i to avoid overlap with the main permutation seeds, and score every column with the active algorithm. From each permutation we retain

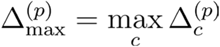

yielding a null distribution of size *P_null_*. For each nominated column c:

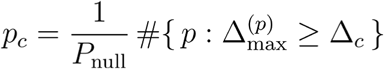

Candidates are walked in descending Δ order. Columns with *p꜀* ≤ 0.05 are removed; the walk breaks at the first failure. If rank 1 fails, pruning halts. Use of max꜀ Δ꜀ controls family-wise error across the alignment without per-column correction. The layer is on by default and disabled with --no-delta-null.

#### Reactive touchdown rollback

A partial batch, fewer than k candidates justified in one iteration, or the alignment-level stop firing, triggers a one-time rollback: the most recent batch removal is undone, the step size is reduced to max(1, k₀ ⁄ 10), alignment and delta-score null are calculated on the pruned alignment, and pruning resumes. Subsequent partial batches are accepted as removed. The loop terminates when rank 1 fails.

### Datasets

#### Simulated datasets

To benchmark performance, we used simulated alignments constructed to exhibit varying levels of compositional heterogeneity and known ground-truth topologies. Dataset design followed the simulation strategies of Foster et al. (2023), including alignments with multiple compositional regimes across the tree. All alignments were simulated with p4 (commit 91e49a9) (Foster 2004).

#### Empirical Null Validation

To approximate a parametric null distribution, we sampled 10 values from a χ² distribution with 19 degrees of freedom (corresponding to 20 amino acid states) in 1,000 iterations using the rchisq() function in R (v4.4.1), generating a total of 10,000 theoretical χ² scores for comparison. To assess the behavior of parametric versus empirical χ² null distributions, we simulated 1,000 alignments under both homogeneous and heterogeneous amino acid composition, across three species trees of increasing branch length (tree length of 0.25, 2.5, and 5.0). Alignments consisted of 10,000 columns and four taxa. Under homogeneous conditions, all taxa evolved under the LG model. Under heterogeneous conditions, *b* branches evolved under the Dayhoff amino acid frequency profile, while the others retained root composition. Simulations were performed using p4 (Foster 2004) commit 70364fb, specifying stationary substitution models and amino acid frequencies directly. These datasets were used to generate observed χ² score distributions for each taxon, as well as to evaluate the accuracy of permutation-based null models and empirical p-values in classifying biased versus unbiased taxa.

#### Pruning Strategy Benchmarking

To benchmark the performance of different pruning strategies, we simulated a separate set of amino acid alignments under a fixed nine-taxon tree topology with two alternative compositions (Dayhoff and GAP-FYMINK). Five of the nine taxa were assigned to evolve under either Dayhoff or GAP-FYMINK compositions, while the remaining four taxa used the root composition. Internal branches were set to 0.01 substitutions per site and external branches to 1.0, creating a challenging scenario with strong compositional signal and shallow clade separations. We simulated individual alignment lengths of 5-, 10-, 20-, and 50-thousand columns. This setup favors artefactual clustering by composition and provides a stringent test of pruning effectiveness.

Pruning was performed with WitChi using three scoring algorithms (squared, quartic, and Wasserstein) and pruning step sizes ranging from 1 column per iteration to 10% of the initial alignment length. All alignments were pruned until alignments were no longer significantly biased or delta-null was no longer significant according to the respective method.

##### Foster et al. (2023) recoding benchmark set

Protein and DNA alignments depicted in Figures 3 and 4 in Foster et al. (2023) with compositional heterogeneity over the tree were downloaded from http://dx.doi.org/10.5061/dryad.6djh9w11s. Pruning was performed with WitChi (https://github.com/stephkoest/witchi/tree/v0.1.0-alpha) using squared algorithm and pruning step size was 1.

##### Timing pruning algorithms

To assess computational scalability, we measured runtime and memory usage for pruning alignments of 10,000 columns while varying the number of taxa, i.e., 10, 50, 100, 200, 1000. Timing benchmarks were performed on a dual-socket AMD EPYC 9654 “Genoa” machine (2 × 96 cores, 2.4 GHz, 384 GiB RAM), running a Linux environment.

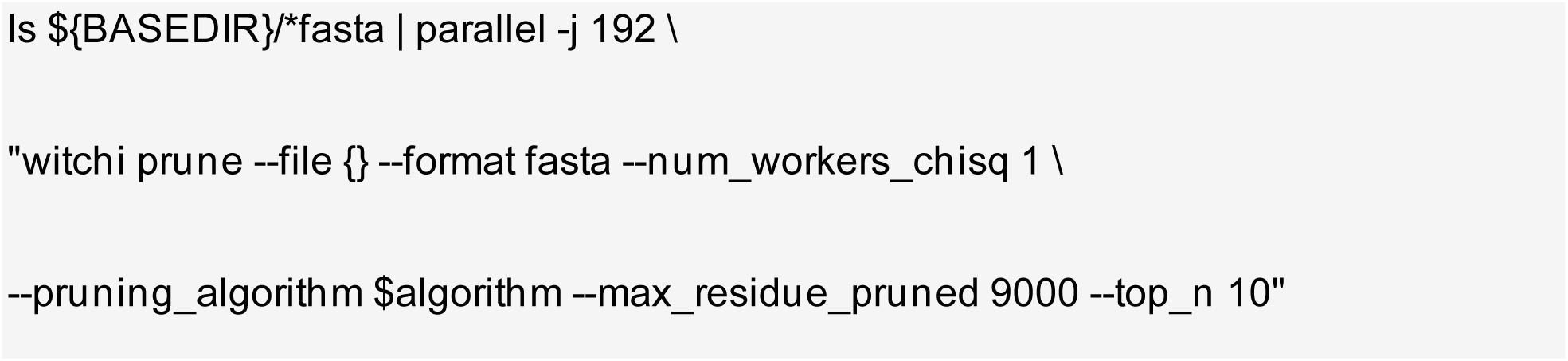

Each pruning run used a single core per alignment (--num_workers 1) to ensure consistent measurement of algorithm scaling with taxon number. Total runtime and memory was recorded for each pruning algorithm (quartic, squared, Wasserstein) across varying taxon counts. BMGE timings were obtained using the stationary-based trimming mode (-s YES) with entropy filtering disabled (-h 0:1), executed similarly in batch mode.

Testing was executed using WitChi’s command-line interface with the following command:

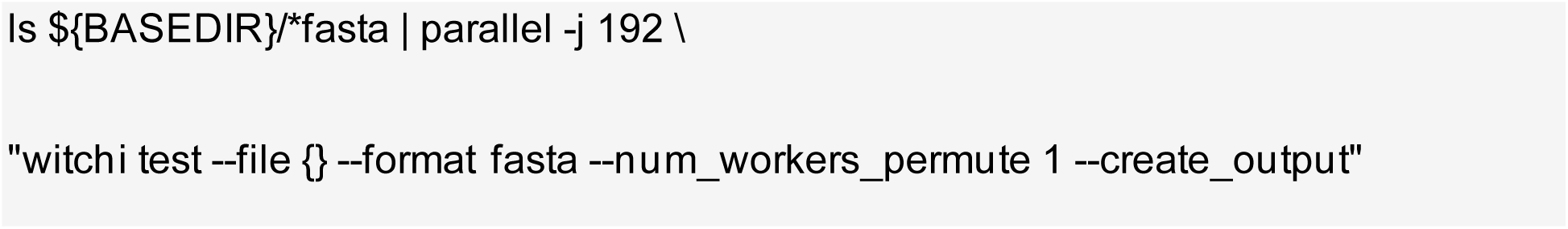

##### Parallel runtime scaling

To evaluate how well WitChi benefits from multi-core execution, we measured runtime of pruning and significance testing operations on alignments of varying size using 1, 2, or 3 CPU cores. For taxon scaling (20–200 taxa), alignment length was fixed at 10,000 residues; for alignment scaling (10,000–100,000 residues), taxon number was fixed at 50. Speedup was calculated as the ratio of runtimes with one core vs. multiple cores (T₁/Tₙ).

#### Estimated overlap calculations

To quantify the similarity between χ² score distributions, we computed the estimated kernel density overlap (eO) between observed scores from homogeneous alignments and two null models (permutation-based and parametric). Kernel density estimates were computed on a common support grid (with a ±1 buffer around the data range) using base R’s density() function. The minimum density at each point (pmin) was integrated numerically using integrate.xy() from the sfsmisc package to calculate the area of intersection. The eO coefficient was computed as:

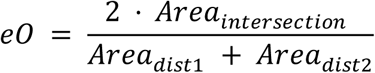

This yields values between 0 and 1, where 1 indicates complete overlap. Overlap was calculated separately for each internal branch length tested (0.25, 2.5, 5.0).

#### BMGE stationary trimming

We benchmarked BMGE v1.12 using its stationary-based trimming mode to compare performance with WitChi. Alignments were trimmed using the following parameters: “-t AA -h 0:1 -s YES”. The -s YES flag enables the stationary-based trimming algorithm, which removes alignment columns based on pairwise composition distance among sequences. To ensure a fair comparison focused solely on composition-based pruning, entropy filtering was disabled by setting the entropy threshold range with -h 0:1. The -t AA option specifies amino acid alignments. All other parameters were left at their defaults. BMGE was applied to the same simulated datasets and evaluated using the same downstream tree inference protocol as used for WitChi.

#### Phylogenetic tree inference from simulated alignments

Unless otherwise specified, all trees from simulated amino acid alignments were inferred with IQ-TREE v2.1.3 (Minh et al. 2020) under the LG+G4 model. DNA alignments were inferred with the GTR+G4 model.

#### Recoding and mixture-model reconstructions

To compare WitChi pruning with established remedies for compositional bias, the nine-taxon biased simulations (50,000 columns) were reconstructed under amino-acid recoding, the four-state SR4 and the Dayhoff-category alphabets, with trees inferred under GTR+G4, and under a site-heterogeneous mixture model (LG+C60+G4 in IQ-TREE with optimized mixture weights, --mixt-opt). All reconstructions used the same alignments and the same branch-length / topology comparison (vs the true and homogeneous-control trees) as the WitChi-pruned trees.

#### Comparison of phylogenetic support across pruning algorithms

To quantify how well each pruning strategy recovers the true topology, we measured true-clade recovery across the 100 replicate alignments for WitChi’s squared, quartic, and Wasserstein algorithms and for BMGE stationary trimming. For each algorithm, alignment length, and pruning step size, recovery is the number of replicates (out of 100) in which the correct clade is present in the inferred best tree. Because shorter alignments carry less information and do not always recover the true tree even under compositional homogeneity, recovery is expressed relative to the unpruned homogeneous control (compositionally homogeneous alignments generated under the same tree), as a percentage of the control’s recovery. We report this recovery relative to control per algorithm × alignment length × step size (Fig. 3D) and summarized across the four lengths at each pruning intensity (Fig. 3E). As a control, pruning the homogeneous alignments themselves recovers the true tree at essentially the control level (dashed lines in Fig. 3D), confirming that pruning a bias-free alignment does not degrade recovery.

#### Benchmark data processing and visualization

All statistical analyses and figures were generated using R v4.4.3 (R Core Team 2018). Data processing utilized data.table v1.16.4 (Barrett et al. 2025). Visualization was performed using ggplot2 v3.5.1 (Wickham 2009), cowplot v1.1.3 (Wilke 2015) for multi-panel figure assembly, and patchwork v1.3.0 (Pedersen 2019) for composite layouts.

#### Empirical Dataset

To evaluate WitChi’s performance on empirical data, we selected the archaeal dataset from GTDB r220 (Rinke et al. 2021). The MSA, consisting of 53 archaeal marker genes (Dombrowski et al. 2020), includes 5,869 taxa and 10,101 columns. We evaluated χ^2^ values with WitChi and pruned the supermatrix using the Wasserstein algorithm, which is the highest-ranking from benchmarking. The pruning was set to a step size of 10 residues per iteration and after the removal of 51.14% of the original alignment none of the remaining columns had a Δ-score that exceeded the homogeneous expectation defined by the Δ-null. FastTree v2.1.11 (-lg) (Price et al. 2010) was used to reconstruct a phylogeny from the pruned supermatrix. The resulting tree served as a guide in a PMSF approximation (Wang et al. 2018) within IQ-TREE v2.1.3 under the LG+C10+G4+F model, with branch support assessed using 1,000 ultrafast bootstraps (UFboot) (Hoang et al. 2018) and 1,000 SH-like approximate likelihood ratio tests (SH-aLRT) (Guindon et al. 2010). To assess patterns of incongruence, we constructed a tanglegram comparing the reference archaeal phylogeny of GTDB with the phylogeny reconstructed from the WitChi-pruned GTDB supermatrix, using the phytools v2.0 R package (Revell 2024). In both trees, leaves of species representatives were collapsed at either phylum or class level. For each group, we retained the species whose terminal branch length was closest to the median branch length of its corresponding phylum or class. The optimal growth temperature for all archaea species representatives of GTDB r220 was estimated using GenomeSPOT v1.0.1 (Barnum et al. 2024).

## Results

### WitChi’s column permutation provides a robust null hypothesis for alignment χ² testing

To evaluate whether WitChi’s empirical null distribution provides a reliable reference for unbiased alignments, we simulated alignments evolving under both homogeneous and heterogeneous composition, and under trees with varying internal branch lengths to modulate divergence. Alignments simulated under homogeneous composition provide the gold-standard reference for expected χ² score distributions given a known species tree (simulation-based null). As expected, observed χ² scores increased with evolutionary distance and were further elevated when taxa evolved under distinct compositions—reflecting the accumulation of compositional signal (Fig. 2, top row).

**Figure 2.**
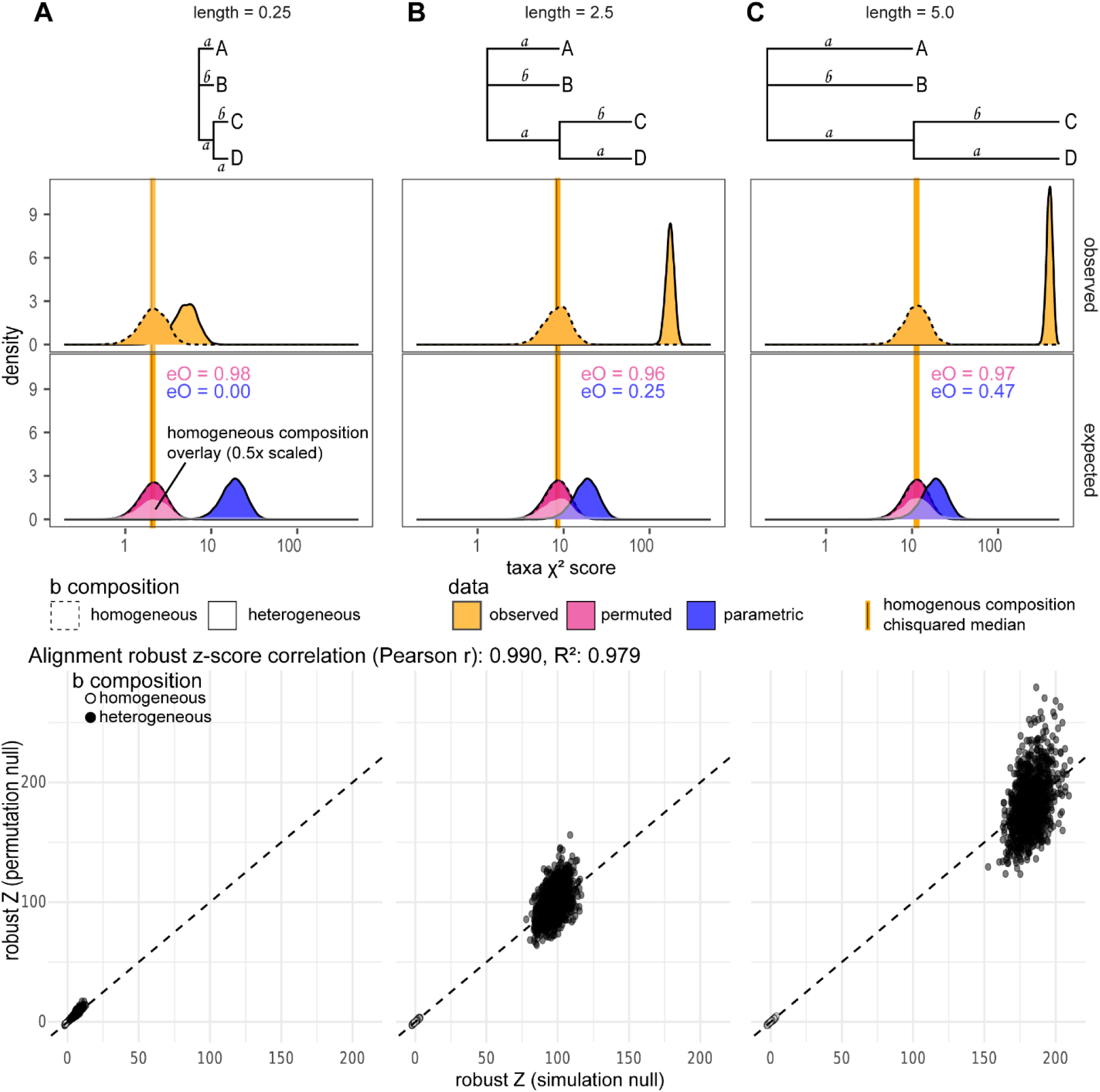
Column permutation of heterogeneous alignments approximates a homogenous null distribution. (A–C) Phylogenetic trees in the first row illustrate three scenarios with varying tree lengths (0.25, 2.5, 5.0). Density plots in the second row show observed taxa χ^2^ score distributions for 1,000 alignments with heterogeneous compositions (solid lines, where branches b evolve under a different amino acid composition) and homogeneous compositions (dashed lines), respectively. The third row density plots compare expected χ^2^ distributions derived from permuted alignments (pink) and parametric χ^2^ distributions (blue). Observed scores are represented by orange distributions, with solid vertical orange lines marking the median χ^2^ score for homogeneous compositions. The panels highlight the increase of χ^2^ scores as total tree length increases, demonstrating the divergence between observed, permuted, and parametric distributions. eO represents the estimated kernel density overlap between the χ^2^ distribution of alignments with homogeneous composition and either the permuted alignment χ^2^ distribution for heterogeneous composition (pink font) or the parametric χ^2^ distribution (blue font). Scatter plots in the bottom row compare alignment-level robust Z-scores from the simulation null (x-axis) and the permutation null (y-axis), pooled across the three tree lengths; the diagonal dashed line indicates a slope of 1.

To approximate a null model without requiring tree inference and simulation, WitChi generates permuted alignments by randomly shuffling characters within each alignment column (Fig. 1). This disrupts taxon-specific composition while preserving site-wise residue distributions, thereby retaining the alignment’s global structure (permutation-based null).

For comparison, we also evaluated the parametric null, i.e., the canonical χ² distribution that assumes independent and identically distributed samples with degrees of freedom equal to the number of character states minus one. Across all conditions, parametric χ² distributions increasingly failed to reflect the expected null distribution, as shown by a decline in kernel density overlap (eO) between the homogeneous taxa χ² scores and the parametric null (Fig. 2). This overlap dropped to near zero even at modest tree lengths (eO = 0.00–0.47). In contrast, permutation-derived null distributions (see *Methods*) closely matched the homogeneous expectation across all divergence levels, with consistently high overlap values (eO = 0.96–0.98; Fig. 2, third row). This demonstrates that permutation provides a robust, alignment-specific null model that adapts to the structure of each dataset. Although Fig. 2 pools the permuted scores for display, the single-alignment null used in practice varies little between datasets: the per-alignment q95 (P = 100) is tightly concentrated (mean ± SD 3.62 ± 0.14, 14.27 ± 0.46, 18.53 ± 0.59 at tree lengths 0.25 / 2.5 / 5.0; CV < 4%) and coincides with the q95 of the observed statistic on the same null data.

To quantify deviations from the null distribution, we implemented robust Z-score and empirical p-value (also referred to as pseudo p-value) metrics for both taxon-specific and alignment-wide χ² scores. Z-scores were computed as standard deviations from the empirical permutation mean. Under homogeneous conditions, taxon-level Z-scores remained centered near zero. In heterogeneous simulations, Z-scores of biased taxa were strongly elevated (Fig. 2, bottom row). At the alignment level, robust Z-scores from the permutation null closely matched those from the simulation null (Pearson r = 0.990, R² = 0.979; Fig. 2, bottom row), confirming that permutation provides a practical surrogate for costly simulation-based testing.

Because per-taxon χ² null distributions can be distorted on unevenly sampled trees (Supplementary Results; Fig. S1), we verified on an uneven four-taxon simulation that the cross-taxon mean and MAD underlying our robust Z-score are nonetheless reproduced by column permutation; per-taxon Z-scores and p-values are therefore reported as diagnostics, while stopping relies on alignment-level criteria. The same permutation principle extends to the per-column level: the largest per-column Δ-score expected under compositional homogeneity is reproduced by column permutation, providing a per-column "delta-null" that flags columns carrying genuine compositional bias (Fig. S2). The per-column delta-null and the alignment-level empirical p-value together define WitChi’s two stopping criteria (Methods).

### WitChi pruning algorithms efficiently remove compositional bias

To assess the effectiveness of WitChi pruning strategies under realistic conditions, we benchmarked performance on simulated alignments with known true topologies. These datasets were designed to introduce controlled compositional heterogeneity, allowing systematic evaluation of how different pruning approaches impact phylogenetic accuracy. Specifically, we simulated 100 alignments under a nine-taxon tree evolving under the LG+G4 model, in which five taxa (5 terminal branches and 1 internal branch) evolved under two compositions distinct from the root, with ten-fold over- and underrepresentation of specific amino acid sets (STNDHMIF-AGEQKLVW –Dayhoff– and GAP-FYMINK) (Fig. 3A). This setup models patterns of lineage-specific compositional bias and allows direct testing of pruning effectiveness in recovering the true tree using standard substitution models in the presence of multiple compositions.

**Figure 3.**
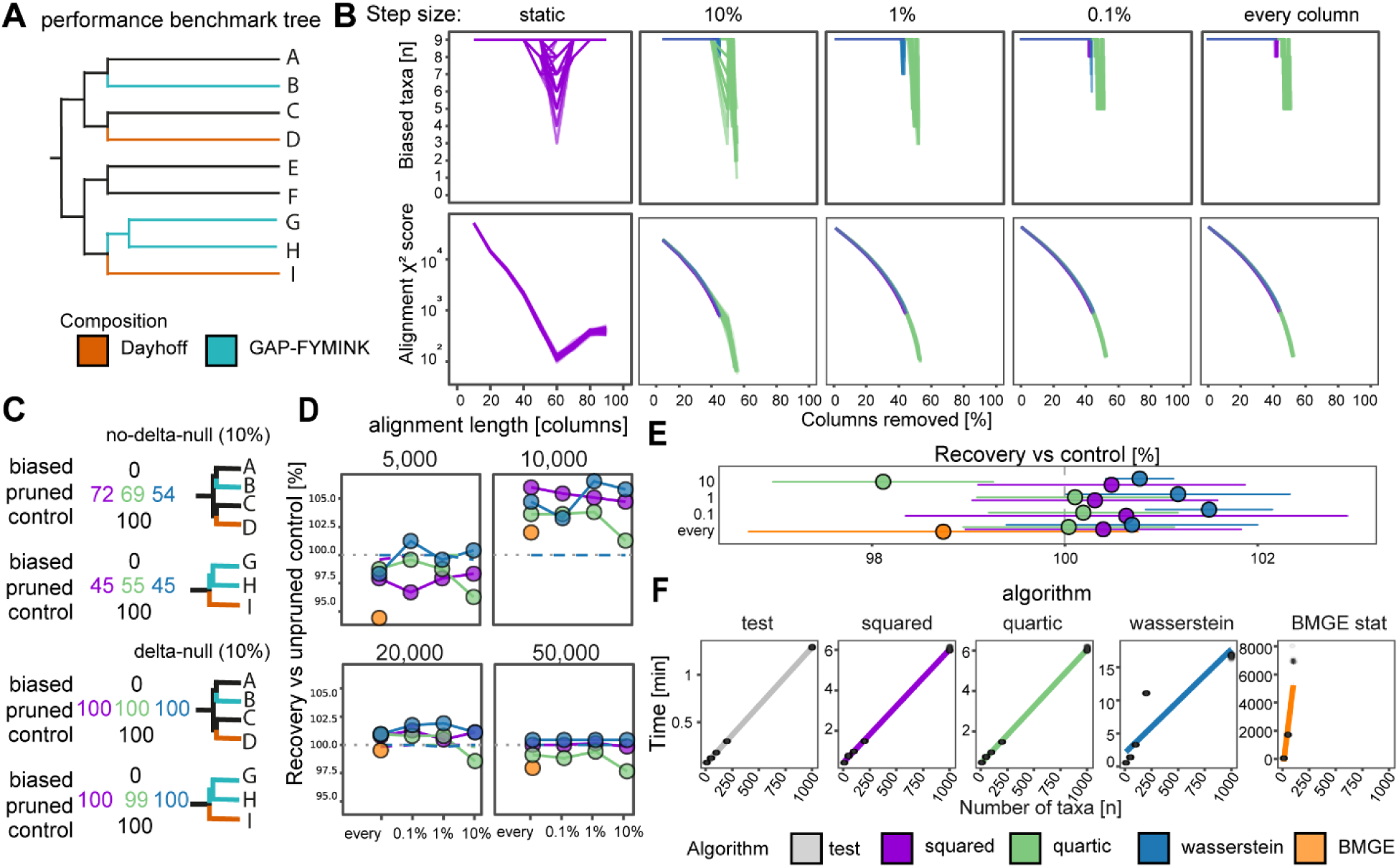
Wasserstein-guided pruning produces the most consistent true clade recovery, and all algorithms scale linearly with the number of taxa.. (A) Tree and compositions used for generating 9 taxa alignments with branches evolving under 3 compositions used for panels B-C: root (black), Dayhoff (orange), and GAP-FYMINK (turquoise). Internal branches were 0.01, and terminal branches 1 substitution per position long. (B) Lines indicate biased taxa and alignment χ² scores when pruning 100 alignments with 50,000 columns using different amino acid frequency recalculation strategies for scoring: no recalculation (static), recalculation after removing every 10%, 1%, 0.1% columns of the initial alignment, and recalculation after each individual column is removed. Line colors represent pruning algorithms (legend: bottom right). (C) Recovery of the true clade from the 50,000 column long simulated alignments from the heterogeneous (biased), the pruned, and the control alignments. The correct clade was almost fully supported when alignments were pruned with step sizes of 10% with –delta-null. (D) True-clade recovery of each pruning strategy (including BMGE stationary trimming) relative to the unpruned homogeneous control, for alignments of 5,000, 10,000, 20,000, and 50,000 columns. Dashed lines show the recovery of the pruned homogeneous-control alignments per algorithm (≈ control level, confirming that pruning a bias-free alignment is safe). See Fig. S3B for detailed per-clade recovery. (E) Each point represents the median recovery relative to control, summarized across the four alignment lengths at each pruning intensity. Whiskers indicate the MAD. (F) Points indicating pruning time of alignments with 10,000 columns and varying number of taxa for significance testing under the three WitChi pruning algorithms, and BMGE stationary trimming.

Recalculating alignment-wide amino-acid frequencies as columns are removed is essential (Fig. 3B). With static global frequencies (no recalculation), bias was never resolved: χ² scores fell at first but rose again after ∼60% of columns had been removed, so static pruning ultimately exacerbates compositional bias. Recalculating frequencies avoided this: across step sizes from single-column removal to 10% of the alignment per iteration, χ² scores fell steeply toward the null and the number of biased taxa decreased, though never to zero. Some taxa stay formally biased because pruning halts at the stopping rules (the per-column Δ-null and the alignment-level empirical p-value) before every taxon is unbiased; this is by design — the Δ-null stops removal once no column exceeds the maximum expected under the permutation null, sparing genuine signal rather than over-pruning. At step sizes of 1% or less the three scorers behaved comparably, removing 44–55% of columns before convergence (≈44% for squared and Wasserstein, ≈52–55% for quartic).

This bias reduction yields accurate topologies, and the gain is robust to step size owing to the Δ-null. Unpruned heterogeneous alignments never recovered the true clades (0%), whereas pruned alignments recovered them at the homogeneous-control level. With the Δ-null enabled (the default), recovery stayed at the control level and the removed fraction was stable (∼44–55%) from single-column steps up to 10% (Fig. S3A); disabling it (--no-delta-null) collapsed both at the 10% step, where pruning over-shot to a median of 74% of columns, with 22–44% of replicates hitting the 90% max-residue cap, and recovery fell to 47–61%. Fig. 3C shows this directly at the 10% step: with the Δ-null the deepest clades (ABCD, GHI) recover at ∼100%, matching the control, whereas without it only 45–72% of replicates recover them. The Δ-null therefore preserves accuracy across step sizes, letting large, fast steps be used without over-pruning.

### WitChi enables true clade recovery under compositional heterogeneity

Having established that dynamic recalculation with the Δ-null restores recovery robustly across step sizes, we next asked whether the three scoring algorithms differ in recovering the true topology (Fig. 3D), testing shorter alignments (5,000–20,000 columns) to probe lower information content. We compared squared, quartic, and Wasserstein across alignment lengths from 5,000 to 50,000 columns (Fig. 3D). Wasserstein-guided pruning recovered the true clades most consistently, ranking first across step sizes and matching or exceeding the homogeneous control, with squared close behind and quartic the weakest of the three.

Pooling across all pruning regimes and alignment lengths, the Wasserstein algorithm with a 0.1% step size gave the highest true-clade recovery relative to the homogeneous control (Fig. 3E; per-clade detail in Fig. S3B), balancing bias reduction and phylogenetic accuracy across alignment lengths.

Pruning remained effective when only part of the alignment was biased (Fig. S4): with bias restricted to 50% of sites, clade recovery rose from 25.9% in the unpruned alignments to 96.3–97.0% after pruning, matching the homogeneous control (97.9%). The three scorers removed columns in essentially the same order and reached the same precision at matched recall (∼0.91), so the delta-null criterion calibrates pruning depth to the realized biased fraction rather than over-trimming the homogeneous remainder.

WitChi pruning also recovered accurate topologies under branch-length heterogeneity (Fig. S6): mean per-clade recovery rose from 42.9% in the unpruned heterogeneous alignments to ∼97–98% across the three scorers, and whole-topology recovery from 0% to 81–84%, approaching the homogeneous ceiling (98.6%). χ²-guided pruning shortens the tree overall (it removes the highest-divergence columns), so the relevant question is whether relative branch-length proportions are preserved. Pruning restored the distance of the internal branches from the true tree to the homogeneous floor (internal branch-score distance 0.009 for squared and Wasserstein, 0.012 for quartic, versus 0.128 for the unpruned alignment; floor 0.009), and Wasserstein also best preserved the terminal proportions (leaf distance 0.080, versus 0.091 squared and 0.133 quartic). On the homogeneous baseline essentially no columns were removed, so branch lengths were unchanged.

We next asked whether WitChi pruning recovers the true topology and branch lengths as well as the established remedies for compositional bias — amino-acid recoding (SR4, Dayhoff4) and a site-heterogeneous mixture model (LG+C60+G4) — on the strongly biased 50,000-column simulations (Fig. S5). On this dataset, standard LG+G4, both recodings, and the LG+C60+G4 mixture all failed to recover the internal topology, each missing four of the six internal clades (AB, CD, ABCD and GHI recovered in ∼0% of replicates), whereas WitChi pruning recovered them as well as the homogeneous control (∼90–100%; zero missing clades). The same held for branch lengths: the distance of the internal branches from the true tree was 0.56–0.71 for the recoded and mixture reconstructions but only 0.17–0.18 for the WitChi-pruned alignments, indistinguishable from the homogeneous control (0.18). On this benchmark, removing the biased columns therefore outperforms recoding or modeling the heterogeneity for recovering both topology and internal branch lengths.

To further validate these findings, we applied WitChi to amino acid and DNA alignments with compositional heterogeneity over the tree from Foster et al. (2023), originally designed to evaluate recoding strategies under strong compositional heterogeneity. In amino acid simulations, pruning with WitChi consistently restored the correct tree in 100% of replicates using a simple LG+F model (Fig. S7). DNA alignments also showed marked improvements under GTR+G4 (Fig. S8). These results confirm that empirical χ²-guided pruning can in principle remove misleading compositional signal, even without recoding or model heterogeneity (Supplementary Results).

### WitChi outperforms alternative pruning strategies and scales linearly

Finally, we assessed computational scalability by measuring runtime for pruning alignments of 10,000 columns while varying the number of taxa (Fig. 3F). All WitChi algorithms scale approximately linearly with taxon number. Wasserstein-guided pruning is the most accurate but also the slowest: because it compares full per-taxon score distributions at each iteration (Fig. 3F), its runtime grows super-linearly with taxon number, reaching ∼17 minutes at 1,000 taxa versus ∼6 minutes for the squared and quartic scorers (which finish in ∼1.5–6 minutes up to 200 taxa). Accuracy and runtime therefore trade off across scorers.

We also benchmarked BMGE’s stationary-based trimming mode for comparison. While BMGE performed comparably to WitChi in true-clade recovery (Fig. S3B), it recovered fewer clades than the best WitChi configurations across alignment lengths (Fig. 3D) and overall (Fig. 3E). Notably, BMGE scaled poorly with increasing taxon count, exceeding 4 days at 100 taxa. We only obtained 80 replicates for the 50 taxa, and 10 for the 100 taxa dataset and we omitted running 200 and 1000 taxa due to a 5-day computation time limit on our available computational resources. BMGE’s stationary trimming approach, while conceptually related, is not optimized for large phylogenomic matrices and scales quadratically with taxon number due to its all-pairs comparison strategy. In contrast, WitChi’s taxon-specific scoring and empirical null framework scale linearly, making it applicable to much larger datasets.

These results support Wasserstein-guided pruning at a relative step size of 0.1% as the default for accurate recovery, with the faster squared and quartic scorers preferable when runtime is the constraint on very large alignments.

Finally, we evaluated parallel runtime scaling using 1–3 CPU cores across varying taxon numbers and alignment lengths (Fig. S9). Pruning operations scaled less efficiently than significance testing, with longer alignments and higher taxon numbers benefiting more from additional cores. The highest speedup was observed for the Wasserstein algorithm, indicating that WitChi particularly benefits from parallelization on large datasets.

### Application to a deeply divergent and compositionally biased archaeal dataset

To assess the effectiveness of WitChi on real data, we applied it to the archaeal supermatrix of GTDB release r220. This dataset provides an ideal benchmark due to its evolutionary significance and known cases of compositional bias affecting deep archaeal relationships, particularly among thermophilic and halophilic clades (Eme et al. 2023; Baker et al. 2024; Huang et al. 2025). Using WitChi’s ‘test’ workflow, 5,583 out of 5,869 taxa (95.1%) had an empirical p-value below 0.05, indicating significant bias from the permuted null distribution. We then pruned the 10,101-column alignment with the Wasserstein algorithm and a 0.1% step size—settings optimized through prior benchmarking. WitChi removed 51.14% of columns in under one hour, reducing the number of biased taxa from 95.1% to 3.16% (186 taxa). Pruning halted automatically once no remaining column’s Δ-score exceeded the magnitude expected under a homogeneous alignment (the per-column delta-null), preventing over-trimming. This safeguard ensures that pruning ends not only when the alignment appears to be unbiased but also when further removal would likely compromise informative phylogenetic signal.

Phylogenetic reconstruction from the pruned alignment, using the same model as GTDB, revealed several topological rearrangements consistent with prior in-depth phylogenomic studies (Fig. 4A). For example, the class JANJXX01—encompassing Panguiarchaeales and Njordarchaeales—comprises members with biased amino acid compositions (Eme et al. 2023) and that are predicted to grow at high temperatures (Fig. S10). Phylogenomic analyses have placed this class either with the thermophilic Korarchaeia (Qu et al. 2023) or within Heimdallarchaeia (Eme et al. 2023; Huang et al. 2025) Here, JANJXX01 is placed within Asgardarchaeota, consistent with previous inferences that account for compositional bias (Eme et al. 2023; Huang et al. 2025). Similarly, DRAE01 and Korarchaeia—both composed of hyperthermophiles (Fig. S10)—were split from this clade and more plausibly repositioned, with DRAE01 placed within Thermoproteota and with Korarchaeia at the base of the clade formed by Thermoproteota and Asgardarchaeota (Fig. S11), in agreement with recent studies (Huang et al. 2025; Tahon et al. 2023).

**Figure 4:**
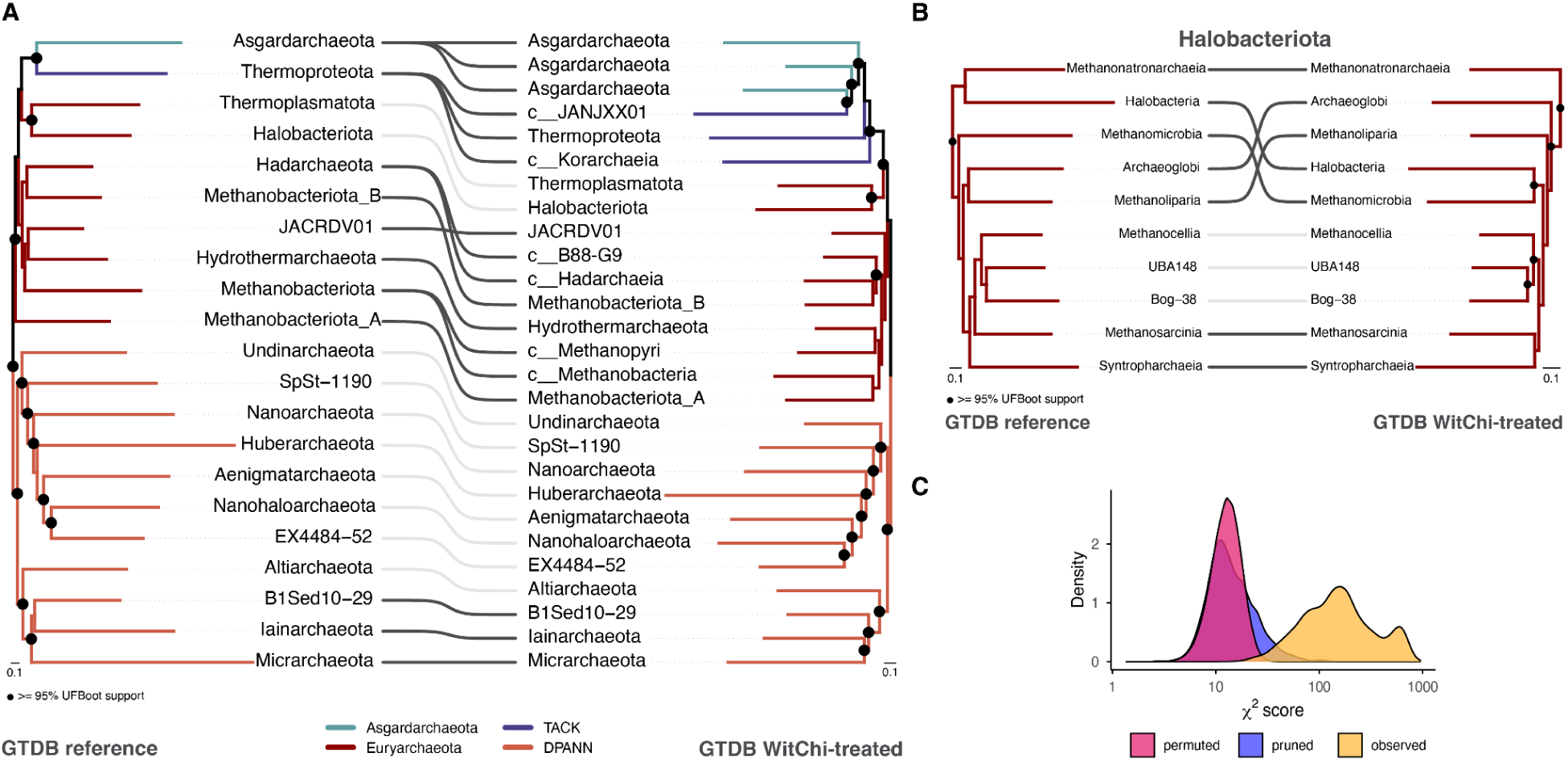
Addressing compositional heterogeneity with WitChi leads to topological rearrangements in the archaeal reference tree of the Genome Taxonomy Database. (A) Phylum-level tanglegram comparing the archaeal Genome Taxonomy Database r220 (GTDB) reference tree (left) with its WitChi-treated counterpart (right). The WitChi-treated phylogeny was inferred from the pruned alignment of 4,935 columns (down from the original 10,101). Light gray links indicate branches with unchanged topology, while colored links highlight topological rearrangements, with each color corresponding to the supergroup of the affected lineage. Phylum-level splits resulting from the WitChi treatment are shown leading to their respective member lineages, with lineages corresponding to single classes labeled with the prefix “*c* ”. Both trees are rooted at the clade shown in orange, originally named after Diapherotrites, Parvarchaeota, Aenigmarchaeota, Nanoarchaeota and Nanohaloarchaeota (DPANN) (Rinke et al. 2013). Black circles at nodes indicate UFboot ≥ 95% support. The scale bar denotes the average number of substitutions per site. (B) Subtrees from (A) focusing on rearrangements in the relationships among the classes within the phylum Halobacteriota. (C) Distributions of χ² scores for the GTDB ar53 supermatrix: pink represents the distribution derived from 100 permuted alignments, orange represents the observed scores in the original alignment, and blue represents the scores for the WitChi-pruned supermatrix.

Another well-documented artifact concerns halophilic archaea. In the GTDB reference tree (Fig. 4B, left), the classes Methanonatronarchaeia and Halobacteria are recovered as sister lineages, suggesting a single origin of halophily. However, this grouping likely reflects convergent compositional bias. In the tree reconstructed from the WitChi-pruned alignment (Fig. 4B, right), Methanonatronarchaeia instead emerges as the sister group to all other Halobacteriota, while Halobacteria is well nested within the phylum—supporting independent adaptation to extreme salinity, consistent with targeted analyses (Baker et al. 2024; Aouad et al. 2019).

Together, these results highlight the capacity of WitChi to resolve multiple sources of bias within a single dataset. By homogenising taxon-specific biases (Fig. 4C) and preserving vertical signal, WitChi recovers more accurate topologies with increased branch support across clades, showing a 21.44% increase in average bipartition support for nodes representing class-level and higher groups after pruning.

## Discussion

Compositional bias is a pervasive source of systematic error in phylogenetic inference (Foster and Hickey 1999; Foster 2004), particularly in deep phylogenomics where lineages may evolve under distinct amino acid or nucleotide compositions due to different selection pressures. Standard substitution models assume compositional stationarity (Goldman 1993), and when this assumption is violated, even otherwise well-fitting models can yield incorrect topologies. While composition-aware models and recoding schemes offer partial solutions (Foster et al. 2023), they remain either computationally expensive, or rigid and insufficiently general for large, complex datasets.

WitChi addresses this challenge by combining classical χ²-based detection of taxon-specific biases with permutation-derived empirical null models. These nulls account for the phylogenetic structure of the alignment by disrupting taxon-specific compositional signal while preserving site-wise composition, enabling more accurate assessment of whether observed taxon scores represent significant compositional deviation. This approach avoids assumptions of taxon independence (Goldman 1993), a statistical oversight that, if not properly addressed, may drive inflated significance and false positives (Copley 2025). By better reflecting the statistical behavior of real-world MSAs, WitChi provides a more robust baseline for identifying true compositional bias.

A key insight from our work is that the combination of empirical permutation-derived null distributions with iterative pruning enables robust elimination of compositionally biased signal, without requiring recoding or complex model fitting. The iterative nature of the algorithm—re-evaluating χ² scores after each batch of columns is removed—allows WitChi to target multiple sources of bias. This strategy, first suggested in the context of χ² pruning by Dharamshi et al. (Dharamshi et al. 2023), is here generalized and formalized into a fully automated and scalable implementation. While several scoring strategies are available, our results indicate that using distribution information (as in the Wasserstein scoring algorithm) improves pruning precision and phylogenetic recovery.

Beyond simulations, WitChi restores accurate topologies in complex empirical datasets using only branch-homogeneous substitution models, as demonstrated using the archaeal GTDB r220 supermatrix. Rearrangements observed in the tree obtained from the WitChi-pruned alignment closely mirror results from studies using recoding or complex profile mixture models, including the repositioning of Panguiarchaeales, Methanonatronarchaeia, and Korarchaeia (Eme et al. 2023; Huang et al. 2025; Baker et al. 2024). That these results can be recovered with a lightweight preprocessing step underscores the utility of targeted, composition-aware pruning.

WitChi also addresses practical concerns of scalability. By relying on matrix operations and parallelization, it completes pruning of alignments with thousands of taxa and tens of thousands of sites in under one hour on a modern workstation. This makes the method broadly applicable to modern phylogenomic studies, where computational overhead often limits the use of complex-model approaches.

While BMGE’s stationary-based trimming mode also aims to reduce compositional artefacts without relying on explicit tree inference, it differs fundamentally from WitChi in both statistical framing and computational design. BMGE applies iterative pruning based on pairwise composition comparisons but lacks taxon-specific significance testing and does not generate an empirical null (Criscuolo and Gribaldo 2010). Our benchmarking showed that BMGE performs well at improving phylogenetic recovery, particularly in smaller datasets, but its quadratic scaling with taxon number and the lack of statistical interpretability limit its applicability in large-scale phylogenomic workflows.

While WitChi is currently implemented for amino acid and DNA alignments, its underlying framework is extensible. Future work could explore its application to codon alignments, or alternative alphabets like recoding schemes or structural alphabets. In some datasets, pruning alone may not resolve all artefacts, but as we show, it substantially reduces misleading signal and enables more accurate reconstruction under tractable models. This is especially important because large datasets can produce deceptively high statistical support for incorrect topologies when model assumptions are violated (Kumar et al. 2012). By removing columns that drive such bias, WitChi helps restore a balance between model fit and statistical confidence, improving not just accuracy but also interpretability of phylogenetic support. Beyond pruning, WitChi serves as a diagnostic tool for identifying taxa with extreme χ² deviations, allowing the underlying biases to be inferred. These taxon-specific compositions are informative for evolutionary models that account for across-branch heterogeneity (e.g., GFmix) (McCarthy et al. 2026). This approach is particularly valuable when excessive removal would mitigate bias but at the expense of phylogenetic signal.

In conclusion, WitChi offers a practical and statistically grounded method for identifying and mitigating compositional bias in MSAs. By coupling empirical null testing with iterative pruning, it enables scalable improvement of phylogenetic inference in datasets where compositional heterogeneity would otherwise distort evolutionary relationships. We anticipate it will be a valuable addition to phylogenomic pipelines, particularly in large dataset, and in studies of early-diverging lineages and other taxa where compositional convergence is common and challenging to model.

## Supporting information

Supplementary_Information_Koestlbacher_WitChi

## Data and code availability

WitChi is available as open-source software under the MIT license at https://github.com/stephkoest/witchi, along with documentation and example workflows. We will provide user support for WitChi for a minimum of two years from the date of publication.

All simulated alignments generated for this study will be made available via Dryad (Reviewer link: https://tinyurl.com/dryadwitchi), and an extended version including, benchmarking processing scripts as well as the processed GTDB archaeal alignment and pruned tree files on the Zenodo Digital Repository (Reviewer link: https://tinyurl.com/zenodowitchi). The reference GTDB r220 archaeal alignment and phylogenetic tree are available from https://gtdb.ecogenomic.org.

## Author contribution

S.K. conceived the study, developed the WitChi software, designed and conducted all benchmarking and simulation-based analyses, and wrote the manuscript. K.P. performed the phylogenomic analysis of the GTDB archaeal dataset and wrote the corresponding Results section. D.T. generated the simulated datasets used for algorithm benchmarking. T.J.G.E. contributed to project discussions and secured funding support for S.K. and K.P. All authors provided critical feedback and helped shape the research, analysis and manuscript.

## Funding

We thank SURF (www.surf.nl) for supporting the use of the National Supercomputer Snellius, facilitated through a grant from the Dutch Research Council (NWO-2021.059). This work was supported by the European Research Council Consolidator and Advanced Grants 817834 and 101142180, respectively (TJGE), the Dutch Research Council VI.C.192.016 (TJGE) and VI.Vidi.243.140 (DT), the Volkswagen Foundation Grant 96725 (TJGE), and the Simons Foundation as part of the Moore-Simons Project on the Origin of the Eukaryotic Cell (Grant 73592LPI; https://doi.org/10.46714/735925LPI) (TJGE).

## Acknowledgments

We thank Peter Foster for sharing simulation code that supported the development and benchmarking of WitChi, and for providing valuable feedback on the manuscript draft. We thank Isabelle Anna Zink, Astrid Marihart, Martin Reitschmied, Manuel Pirker-Ihl, and Bela Hausmann for discussions and feedback during the development of WitChi. During the preparation of this manuscript, the authors used a large language model (Claude, Anthropic) to assist with language editing and with coding tasks such as debugging and refactoring. The conception, design, and implementation of the methods, and all scientific conclusions, are the authors’ own. The authors reviewed and verified all resulting text and code and take full responsibility for the content of this work.

## Notes

### Competing Interest Statement

The authors have declared no competing interest.

### Summary of Updates

The stopping rule has been redesigned. Pruning is no longer halted using per-taxon significance counts, which are sensitive to tree shape; it now stops on the logical OR of two robust criteria: a per-column delta-null (no remaining column's score exceeds the magnitude expected under a homogeneous, column-permuted alignment) and an alignment-level empirical p-value above 0.05. Per-taxon scores are now reported only as diagnostics. Per-taxon deviation is standardized with a robust z-score centered on the cross-taxon mean and scaled by the median absolute deviation. The strict mode and the similarity-stratified permutation strategy have been removed. The recommended default scoring algorithm is now the Wasserstein scorer (1-Wasserstein distance in robust z-score space), based on simulation benchmarks and applied to the empirical GTDB analysis; the previous version recommended the quartic scorer. Across alignment lengths Wasserstein gives the most consistent clade recovery. New analyses validate the permutation null against an independent simulation null at the alignment, taxon, and column levels; test pruning when only part of the alignment is biased and under branch-length heterogeneity; report the effect of pruning on branch lengths; compare WitChi against amino acid recoding (SR4, Dayhoff4) and a profile mixture model (LG+C60+G4); and add memory and runtime scaling and expanded test coverage. Supplementary figures and tables were updated accordingly.

