## Supplementary_Information_Koestlbacher_WitChi for "WitChi: Efficient Detection and Pruning of Compositional Bias in Phylogenomic Alignments Using Empirical Chi-Squared Testing"

### Supplementary results

**Per-taxon empirical null, classification performance, and robustness to tree shape**

At the taxon level, robust Z-scores from the permutation null closely matched the simulation null on balanced trees (Pearson r = 0.997, $R^2$ = 0.994; Fig. S1A). Permutation-based empirical p-values accurately distinguished biased from unbiased taxa relative to the simulation null: across all simulations balanced accuracy was 0.988 (sensitivity 0.985; specificity 0.992), and even under the most challenging short-tree condition, where homogeneous and heterogeneous distributions overlap (Fig. 2A), balanced accuracy remained 0.966 (sensitivity 0.948; specificity 0.985). On an unevenly sampled four-taxon tree (three closely related taxa and one distant outgroup), the individual per-taxon $\chi^2$ nulls are distorted (per-taxon kernel-density overlap eO 0.23–0.48), yet the cross-taxon mean and MAD — the location and scale statistics on which the robust Z-score is built — are reproduced by column permutation (eO 0.82 and 0.97), whereas the median and standard deviation are not (eO 0.28 and 0.18; Fig. S1B). Taxon Z-scores from the permutation null therefore remain in close agreement with the simulation null even on the uneven tree (Pearson r = 0.986; Fig. S1C). Finally, the per-taxon permutation threshold is stable with permutation count: the 95th-percentile threshold from P = 100 permutations differs from P = 10,000 by a median of 2.2% (Fig. S1D), supporting the default of 100 permutations.

**Application of WitChi to Simulated Datasets from Foster et al. (2023)**

To further evaluate the performance of WitChi, we applied our pruning approach to amino acid and DNA alignments simulated under compositional heterogeneity over the tree, as described in Foster et al. (2023) and depicted in their manuscript in Figures 3 and 4. These datasets were originally designed to benchmark the effectiveness of amino acid recoding strategies against compositional biases that mislead phylogenetic inference. In our analyses, we pruned the alignments using WitChi's $\chi^2$ score-guided strategy with the squared algorithm, removing columns iteratively in 0.1% steps until no taxa showed significant bias relative to the permutation-based empirical null model. Subsequent phylogenetic inference was performed using the LG+G4 substitution model, a single-matrix site-homogeneous model that does not account for compositional heterogeneity. This choice provided a conservative test of whether bias reduction through pruning alone could restore phylogenetic signal, without invoking more complex heterogeneous models such as CAT or NDCH.

Across all tested simulation scenarios, WitChi pruning perfectly restored the recovery of the true tree topology (Fig. S7). In alignments characterized by tree-heterogeneous compositions, pruning consistently increased the proportion of correct trees relative to untreated data. Importantly, while Foster et al. (2023) found that recoding could variably increase or decrease phylogenetic accuracy depending on the type of heterogeneity, WitChi pruning alone was sufficient to achieve high recovery rates without modifying the amino acid alphabet.

In simulations with moderate to strong compositional heterogeneity (branch c lengths of

0.05, 0.10, or 0.20 substitutions/site), untreated alignments frequently recovered incorrect topologies consistent with compositional attraction. In contrast, pruned alignments nearly always recovered the correct tree, indicating effective bias mitigation. Notably, this result was achieved while using a standard homogeneous model for tree inference, underscoring that effective removal of biased signal can, in some cases, obviate the need for more complex modeling.

Pruning also showed robust performance in simulations based on heterogeneous DNA compositions (Fig. S8), similarly improving recovery rates across all compositionally biased scenarios using a single-matrix model (GTR+G4).

Overall, these results demonstrate that WitChi pruning is a highly effective strategy for correcting compositional biases in both amino acid and nucleotide alignments, achieving substantial improvements in phylogenetic accuracy without the need for recoding or the application of computationally intensive heterogeneous models.

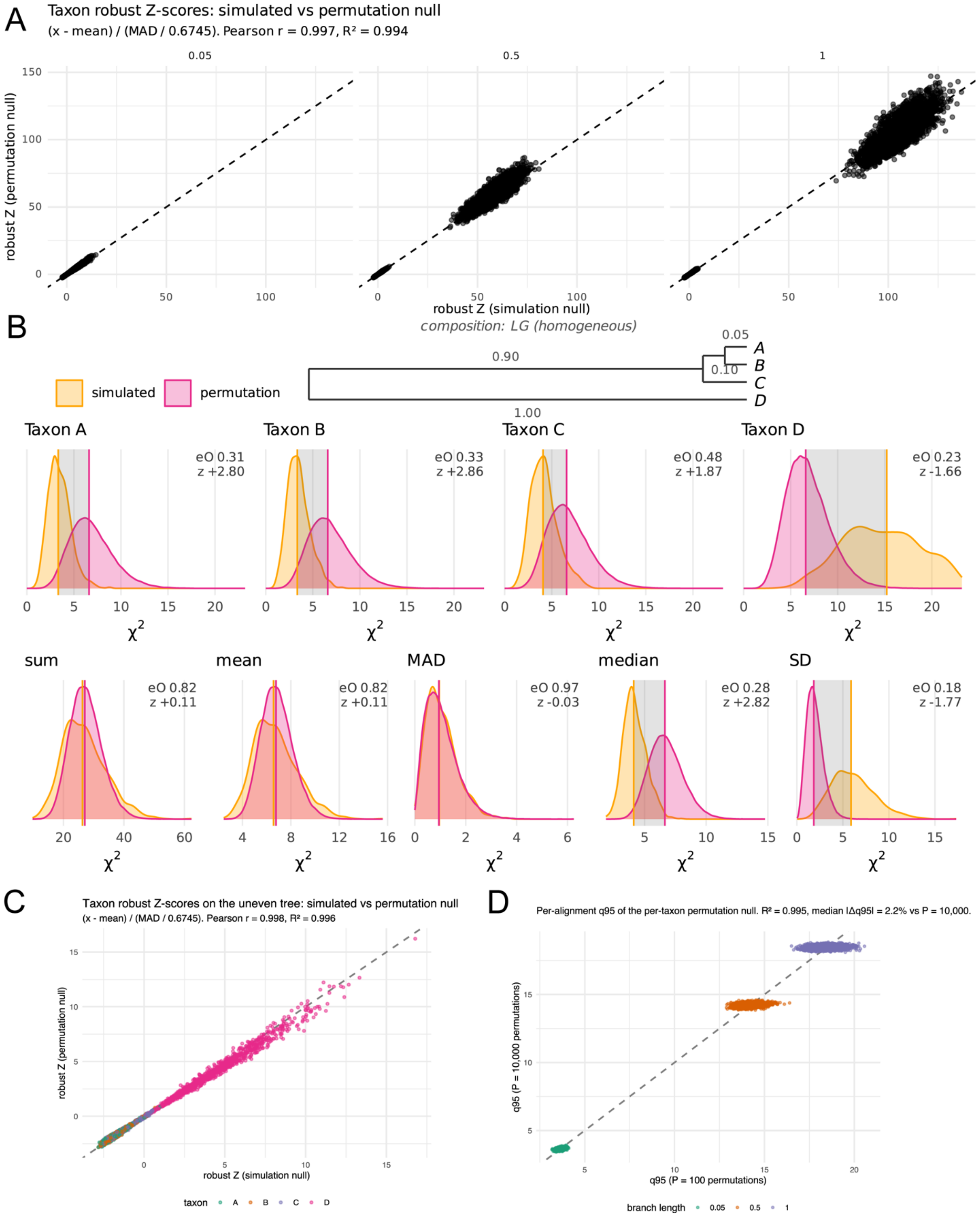


**Figure S1. The robust per-taxon Z-score is reproduced by column permutation regardless of tree shape.** (A) Per-taxon robust Z-scores from the permutation null (y-axis) against the simulation null (x-axis) on balanced trees, one panel per tree length. (B)

The uneven four-taxon tree used as a stress test (three closely related taxa and one distant taxon), showing the per-taxon $\chi^2$ null distributions and the cross-taxon location and scale statistics (mean and median absolute deviation, median and standard deviation) under the simulation and permutation nulls. (C) Per-taxon robust Z-scores from the permutation null against the simulation null on the uneven tree. (D) The per-alignment 95th-percentile permutation threshold computed from 100 versus 10,000 permutations, colored by branch length. Diagonal dashed lines indicate a slope of 1.

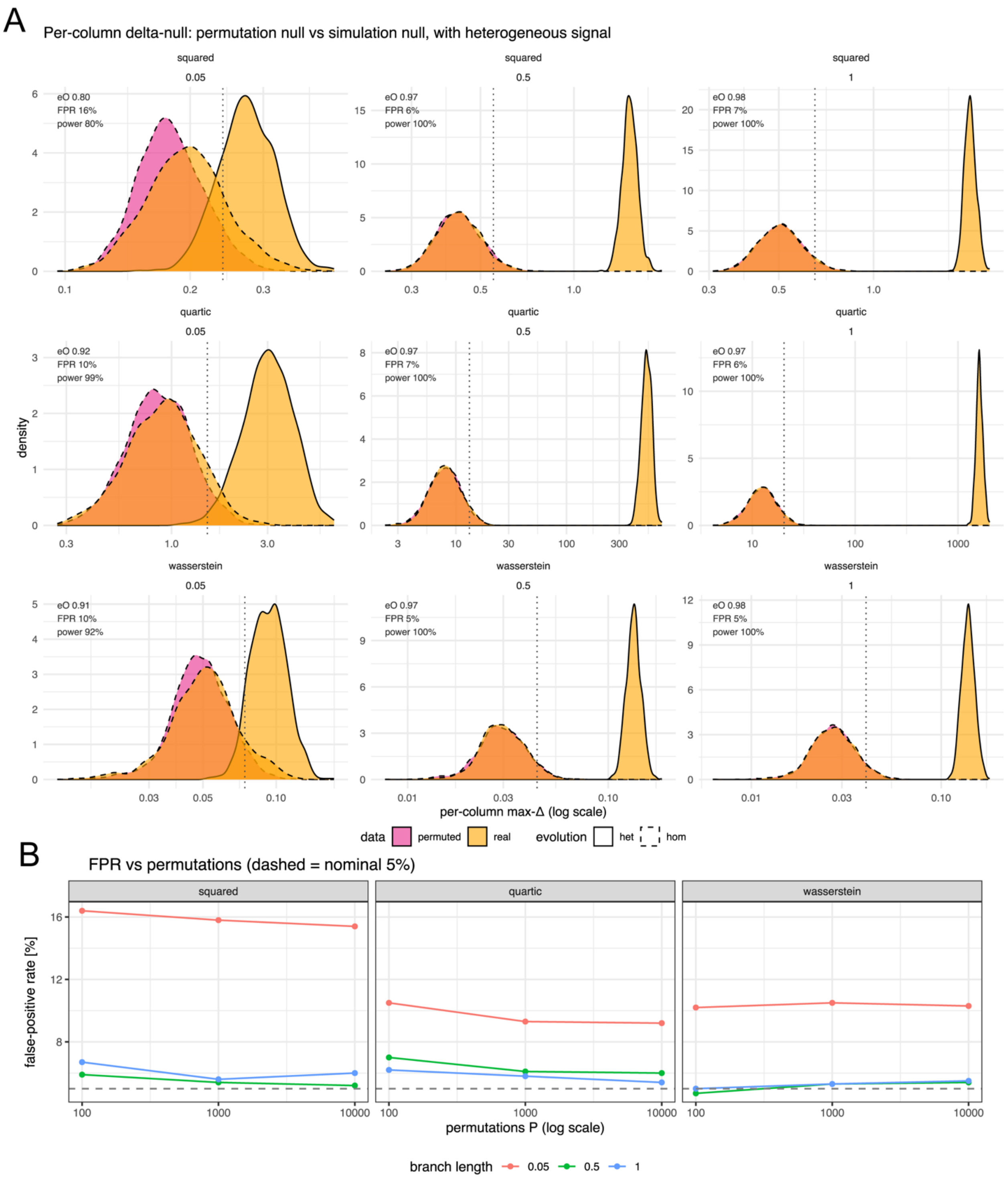


**Figure S2. The per-column delta-null reproduces the column-score magnitude expected under compositional homogeneity.** (A) Densities of the maximum per-column Δ score (raw Δ, log scale) for each scoring algorithm (squared, quartic, Wasserstein) and internal branch length (0.05, 0.5, 1.0). Three distributions are overlaid in each panel: the simulation (ground-truth) null from homogeneous alignments (orange, dashed), the permutation null that defines the stopping criterion (pink, dashed), and the observed scores from heterogeneous alignments (orange, solid); the dotted vertical line marks the median permutation 95th-percentile stop threshold. (B) The fraction of homogeneous alignments whose maximum Δ exceeds the permutation 95th percentile (False-positive rate) as a function of permutation count P (100, 1000, 10,000) for each scoring algorithm and branch length; the dashed line marks the nominal 5% rate.

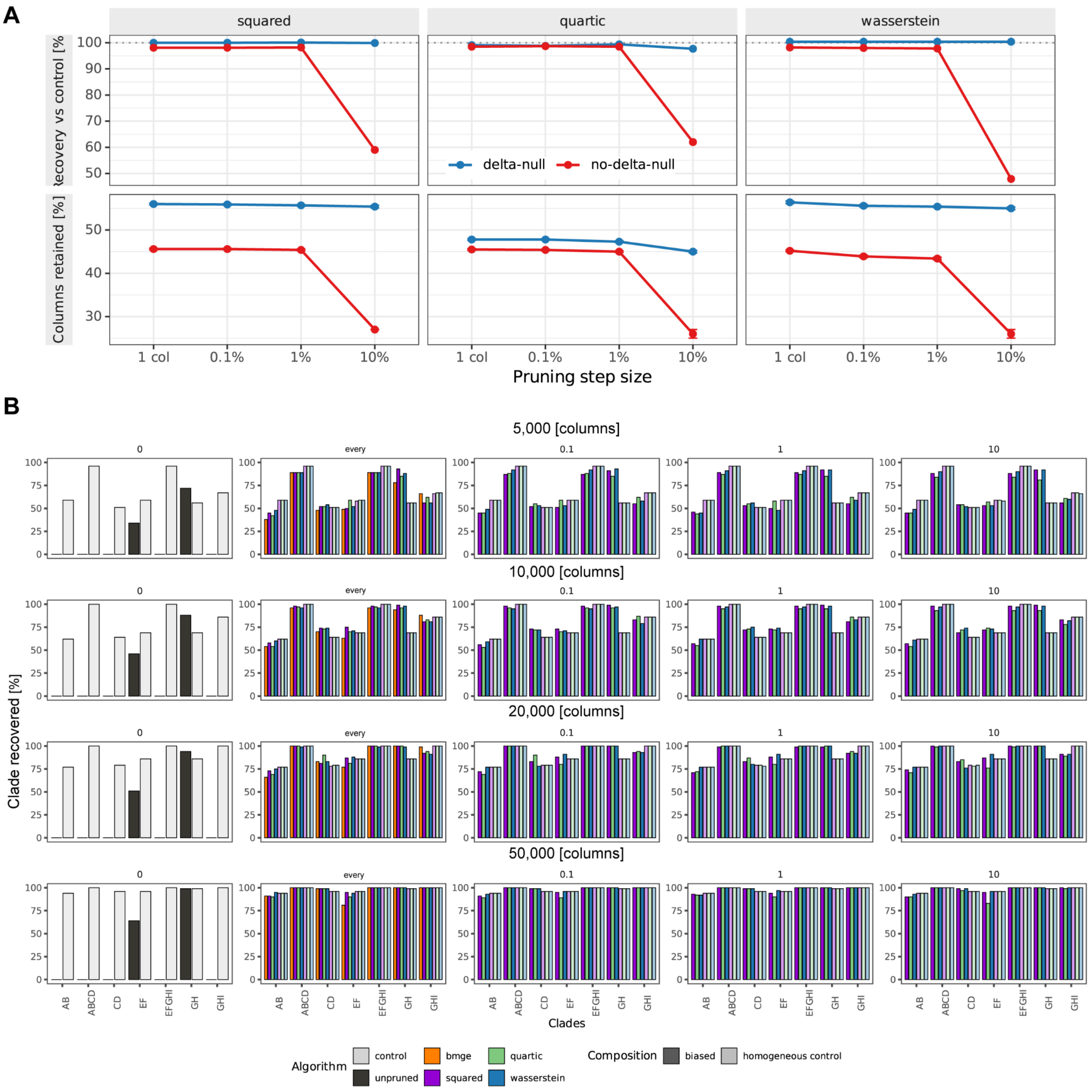


**Figure S3. The per-column delta-null makes pruning depth and topological accuracy robust to the pruning step size.** (A) Nine-taxon alignments simulated under compositional heterogeneity over the tree (50,000 columns, 100 replicates) were pruned with each scoring algorithm (columns: squared, quartic, Wasserstein) at four step sizes (1 column, 0.1%, 1%, and 10% of the alignment removed per iteration), with the per-column delta-null stopping criterion enabled (default, blue) or disabled (--no-delta-null, red). Top row: true-clade recovery relative to the unpruned homogeneous control (total clades recovered as a percentage of the control; the dotted line marks the control, and values may exceed it). Bottom row: percentage of columns retained (median ± median absolute deviation across replicates). With the delta-null enabled, recovery stays at the control level and a stable fraction of columns is retained at every step size; disabling it leaves both near the enabled level until the coarsest (10%) step, where recovery collapses and the alignment is over-pruned — in a fraction of replicates as far as the 90% max_residue cap. (B) Detailed clade-level recovery across pruning strategies and alignment. Bar plots show the proportion of 100 replicate trees that recover each target clade, evaluated across alignments of varying length (5,000 to 50,000 columns; rows) and pruning step size (0%, every, 0.1%, 1%, and 10%; columns). Results are shown for unpruned alignments (black), control alignments simulated under homogeneous composition (light gray), and alignments pruned using the squared (purple), quartic (green), and Wasserstein (blue) algorithms, and BMGE stationary trimming (orange). Each panel includes eight clades of the true tree (AB to GHI) used to quantify recovery accuracy.

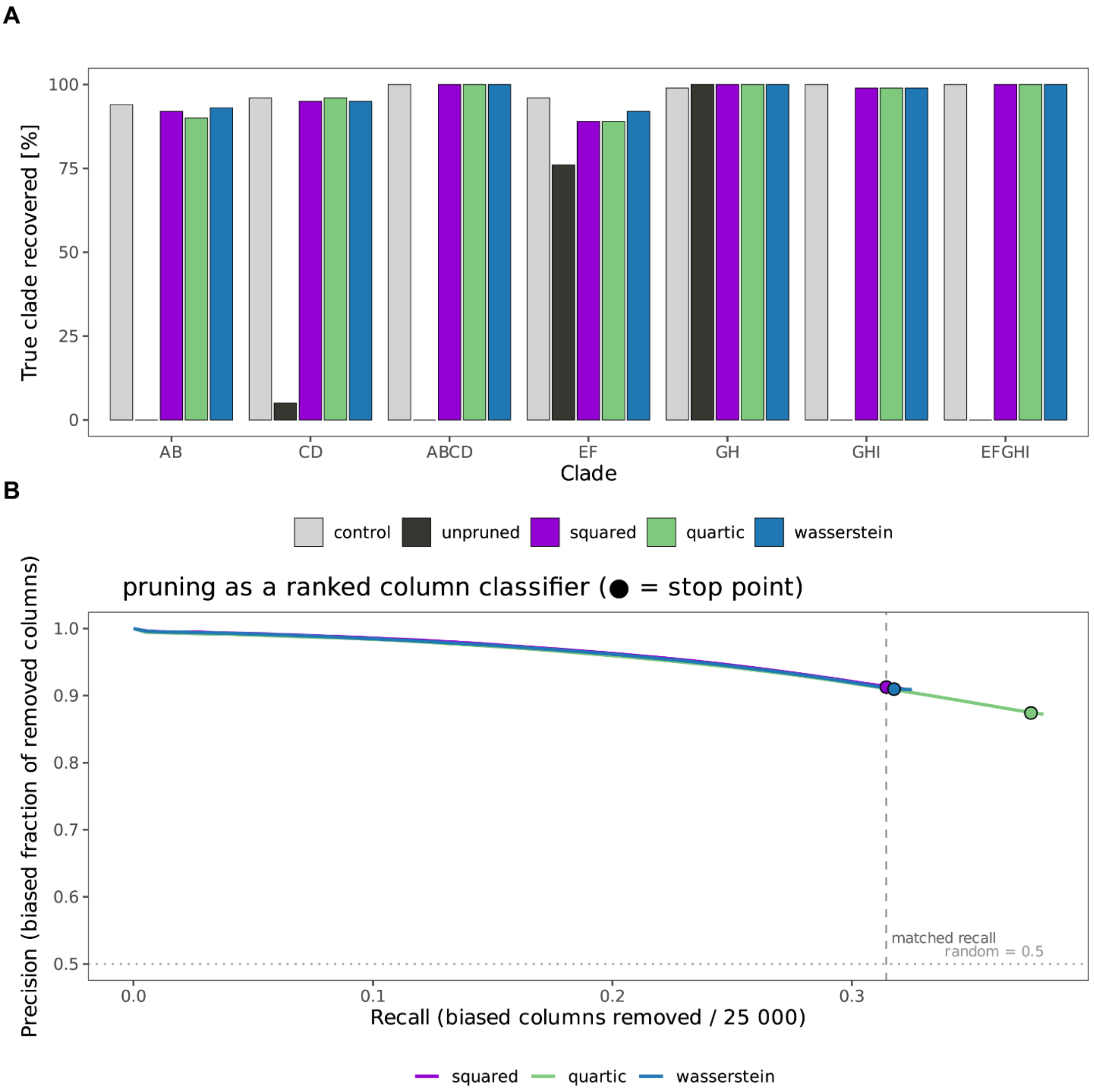


**Figure S4. WitChi pruning recovers the true topology when only 50% of columns in the alignment are affected by compositionally bias.** Nine-taxon alignments were simulated with compositional bias restricted to 50% of sites (the remainder evolving under the root composition; 25,000 of 50,000 columns biased). (A) True-clade recovery (percentage of replicate trees recovering each clade) for the unpruned biased alignments, the alignments pruned with each scoring algorithm, and the homogeneous control. (B) Pruning evaluated as a ranked binary classifier of the truly biased columns, plotting precision (biased fraction of the columns removed so far) against recall (biased columns removed out of 25,000); the filled point marks the stopping point.

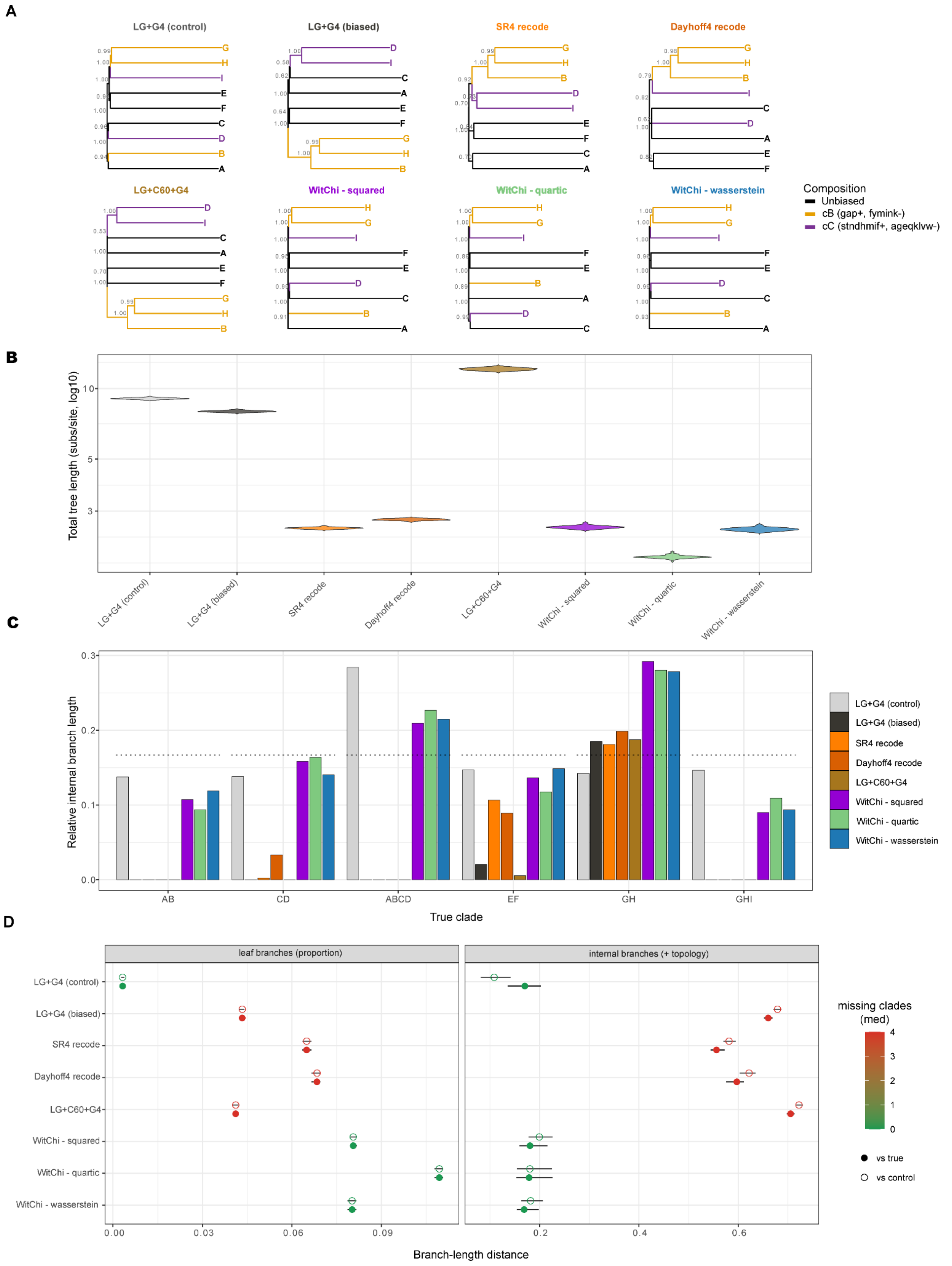
A
LG+G4 (control)
LG+G4 (biased)
SR4 recode
Dayhoff4 recode
LG+C60+G4
WitChi - squared
WitChi - quartic
WitChi - wasserstein
Composition
Unbiased
cB (gap+, fymink-)
cC (stndhmif+, ageqklvw-)
B
Total tree length (subs/site, log10)
C
Relative internal branch length
True clade
AB
CD
ABCD
EF
GH
GHI
D
leaf branches (proportion)
internal branches (+ topology)
Branch-length distance
missing clades (med)
vs true
vs control

**Figure S5. Effect of WitChi pruning, recoding and heterogeneous model on branch lengths.** (A) Consensus trees from 100 replicate reconstructions inferred from the nine-taxon alignments simulated under compositional heterogeneity over the tree (the same scheme as Fig. 3A). Alignments are either used unpruned for tree reconstruction under a homogeneous (LG+G4) or heterogeneous (LG+C60+G4) model, or subjected to SR4 or Dayhoff4 recoding followed by GTR+G4 reconstruction, or pruned with the WitChi algorithms prior to LG+G4 reconstruction; a homogeneous-composition control is shown for reference. Branches are colored by the simulated composition, and node labels give the proportion of replicates recovering each split; trees are rooted and ladderized for display. (B) Total tree length of each consensus tree (log scale); the four-state SR4 and Dayhoff4 recodings are on a reduced substitution scale and are not directly comparable to the amino-acid models. (C) Relative internal branch lengths — each internal branch expressed as a fraction of the summed internal branch length, per true clade — so that differences in overall tree length do not dominate the comparison; the generating (true) tree is shown for reference. (D) Distance of each reconstruction's branch lengths from the true tree and from the homogeneous control, computed separately for terminal (leaf) and internal branches (each normalized to sum to one within its group, then compared by Euclidean distance with union zero-fill so that missing clades are penalized); points are colored by the median number of missing clades.

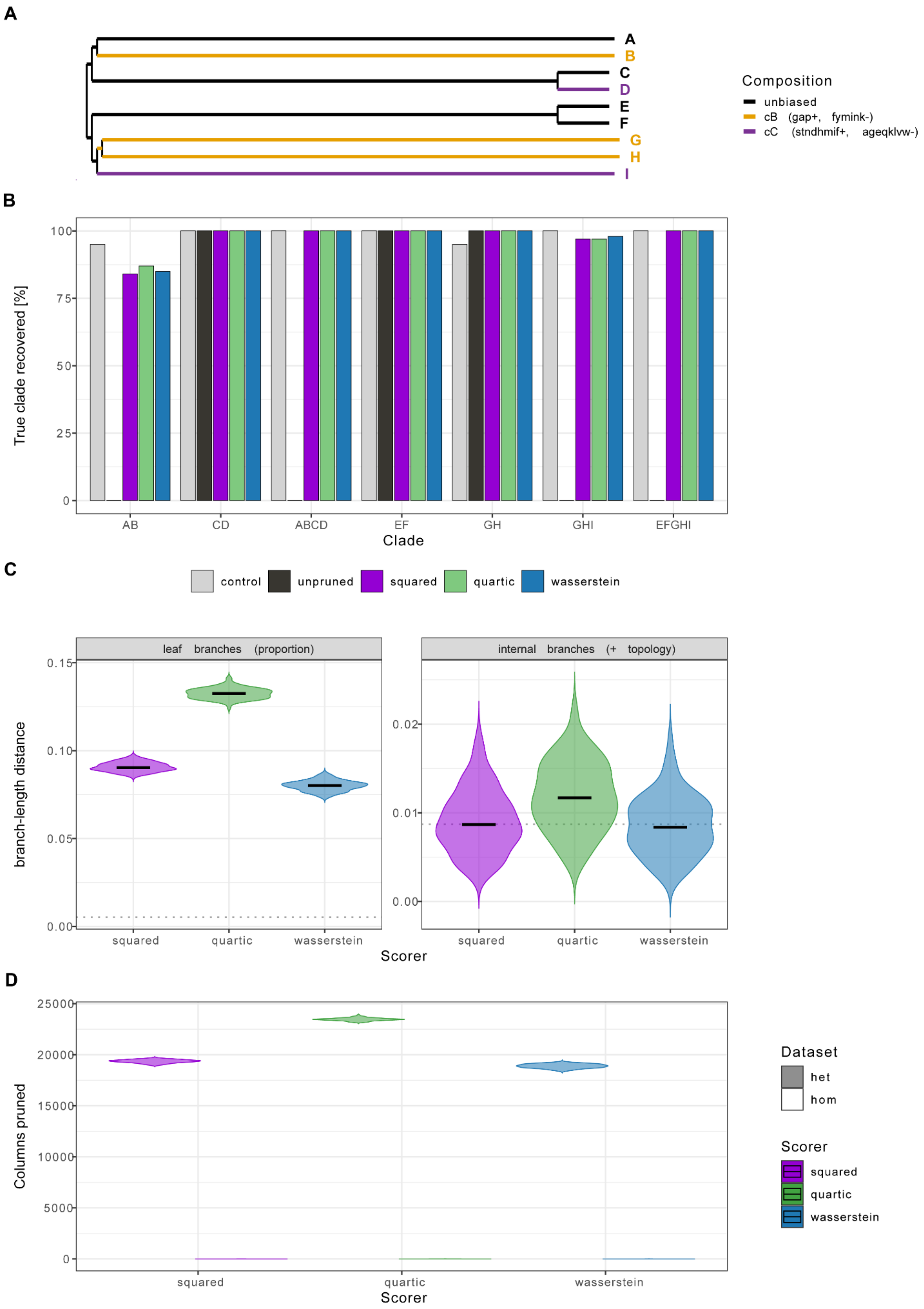
A
A
B
C
D
E
F
G
H
I
Composition
unbiased
cB (gap+, fymink-)
cC (stndhmif+, ageqklvw-)
B
True clade recovered [%]
100
75
50
25
0
AB
CD
ABCD
EF
GH
GHI
EFGHI
Clade
C
control
unpruned
squared
quartic
wasserstein
leaf branches (proportion)
internal branches (+ topology)
branch-length distance
0.15
0.10
0.05
0.00
0.02
0.01
0.00
squared
quartic
wasserstein
Scorer
D
Columns pruned
25000
20000
15000
10000
5000
0
squared
quartic
wasserstein
Scorer
Dataset
het
hom
Scorer
squared
quartic
wasserstein

147

**Figure S6. WitChi pruning recovers accurate topologies under branch-length heterogeneity while preserving relative branch proportions.** (A) Nine-taxon alignments were simulated under the same composition scheme but with branch-length heterogeneity imposed on the CD and EF clades (long terminal stems with tight internal pairs), together with a homogeneous-composition control; pruning used a 0.1% step (100 replicates). The cartoon (top) shows the simulation topology and the lengthened CD and EF stems. (B) Per-clade true-tree recovery for the unpruned heterogeneous alignments, the alignments pruned with each scoring algorithm, and the homogeneous control. (C) Relative branch-length conservation, measured as the branch-score distance on relative branch lengths between the pruned heterogeneous and homogeneous reconstructions, for each scoring algorithm. (D) Pruning depth (number of columns removed out of 50,000) for each scoring algorithm, with the homogeneous control shown for comparison.

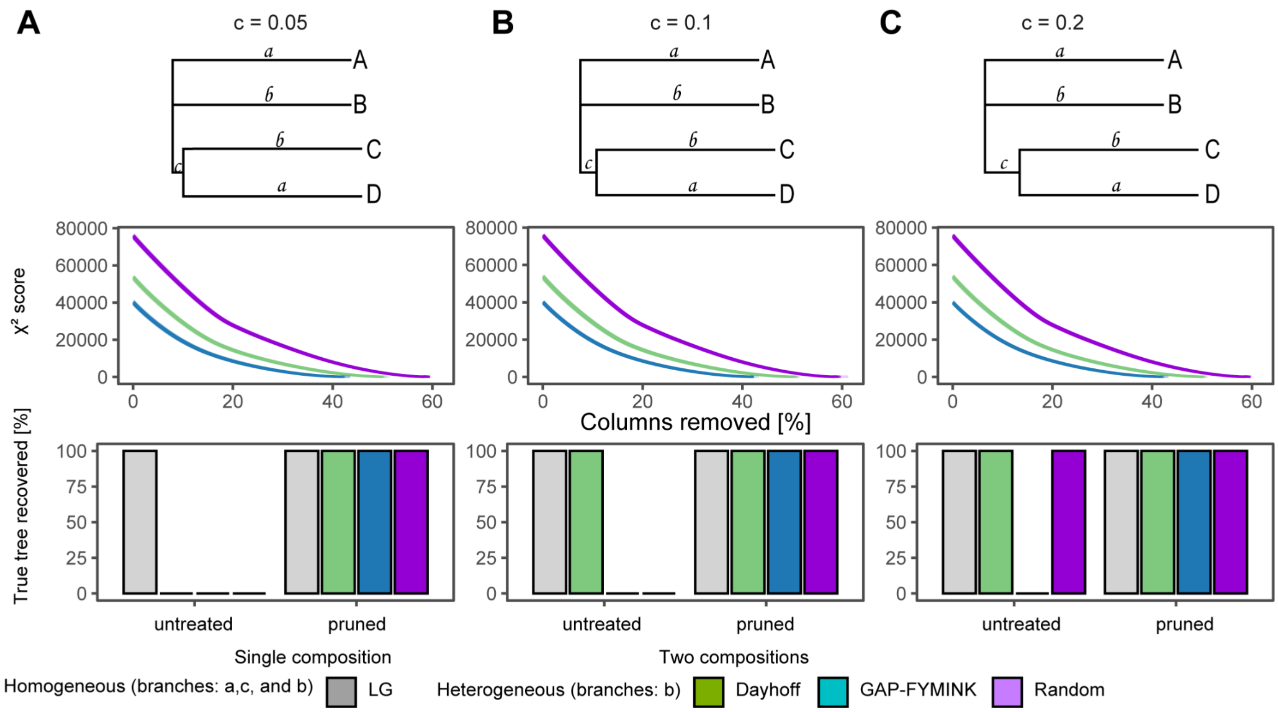


**Figure S7. $\chi^2$ score-guided pruning improves phylogenetic accuracy in amino-acid alignments simulated under heterogeneous compositions over the tree.** (A–C) Phylogenetic trees illustrate three scenarios with varying lengths of branch c (0.05, 0.1, 0.2). Plots in the top row depict the observed $\chi^2$ scores across alignments with distinct heterogeneous compositions (green, blue, purple), as columns are progressively removed. The bottom row shows how often the true tree can be recovered from the untreated (left) or pruned (right) simulated alignments with homogeneous (grey) or heterogeneous compositions (green, blue, purple). Alignments were obtained from (Foster et al. 2023). Pruning removed alignment columns in a 0.1% stepwise fashion until no biased taxa were left.

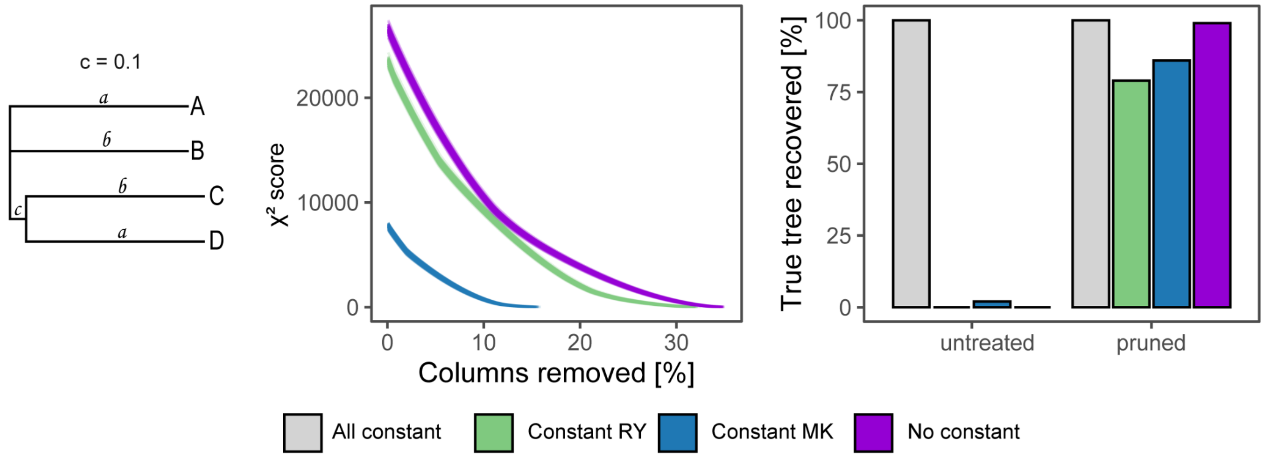


**Figure S8: $\chi^2$ score guided pruning improves phylogenetic accuracy in DNA alignments simulated under heterogeneous compositions over the tree.** Phylogenetic tree under which alignments were simulated. Points in the middle panel show observed $\chi^2$ scores for homogeneous (grey) and heterogeneous compositions (green, blue, purple) during the pruning process. The bar plot shows how often the true tree can be recovered from the untreated or pruned simulated alignments.

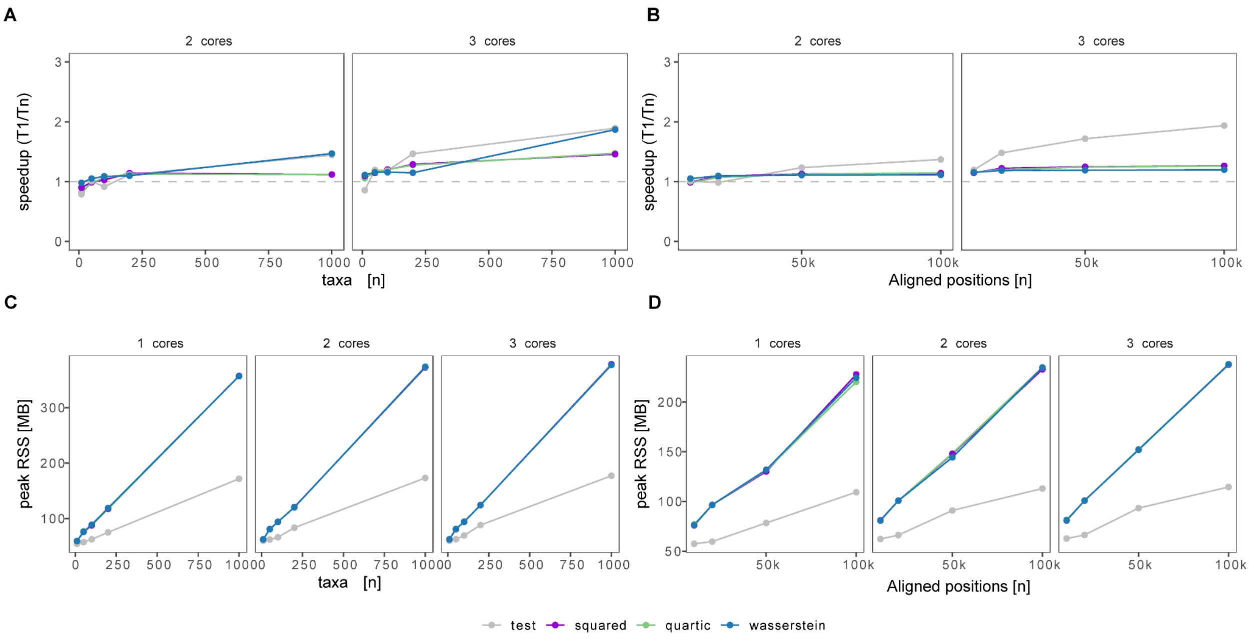


**Figure S9. Parallel runtime and memory scaling of WitChi as a function of taxon number and alignment length.** (A,C) Speedup and peak memory across different taxon counts (20–200) for four WitChi modules (test, squared, quartic, wasserstein) using 1, 2, or 3 CPU cores. (B,D) Speedup and peak memory across increasing alignment lengths (10,000–100,000 positions) at fixed taxon number (n = 50). Speedup is calculated as the ratio between median runtimes with 1 core and *n* cores ($T_1/T_n$). Dashed horizontal lines at Speedup = 1 indicate equal runtimes for multiple cores and a single core.

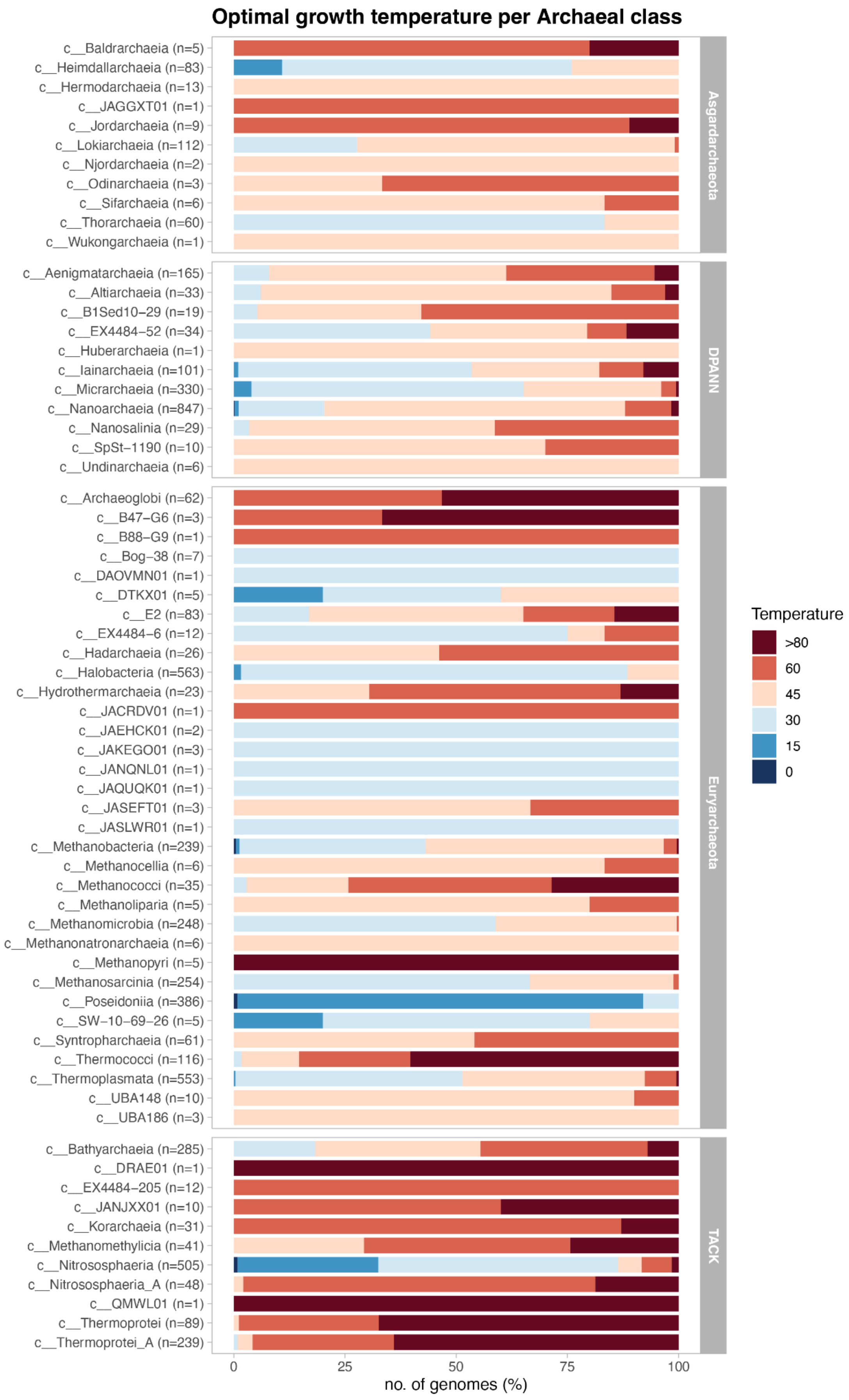

Optimal growth temperature per Archaeal class
c__Baldrarchaeia (n=5)
c__Heimdallarchaeia (n=83)
c__Hermodarchaeia (n=13)
c__JAGGXT01 (n=1)
c__Jordarchaeia (n=9)
c__Lokiarchaeia (n=112)
c__Njordarchaeia (n=2)
c__Odinarchaeia (n=3)
c__Sifarchaeia (n=6)
c__Thorarchaeia (n=60)
c__Wukongarchaeia (n=1)
Asgardarchaeota
c__Aenigmatarchaeia (n=165)
c__Altiarchaeia (n=33)
c__B1Sed10−29 (n=19)
c__EX4484−52 (n=34)
c__Huberarchaeia (n=1)
c__Iainarchaeia (n=101)
c__Micrarchaeia (n=330)
c__Nanoarchaeia (n=847)
c__Nanosalinia (n=29)
c__SpSt−1190 (n=10)
c__Undinarchaeia (n=6)
DPANN
c__Archaeoglobi (n=62)
c__B47−G6 (n=3)
c__B88−G9 (n=1)
c__Bog−38 (n=7)
c__DAOVMN01 (n=1)
c__DTKX01 (n=5)
c__E2 (n=83)
c__EX4484−6 (n=12)
c__Hadarchaeia (n=26)
c__Halobacteria (n=563)
c__Hydrothermarchaeia (n=23)
c__JACRDV01 (n=1)
c__JAEHCK01 (n=2)
c__JAKEGO01 (n=3)
c__JANQNL01 (n=1)
c__JAQUQK01 (n=1)
c__JASEFT01 (n=3)
c__JASLWR01 (n=1)
c__Methanobacteria (n=239)
c__Methanocellia (n=6)
c__Methanococci (n=35)
c__Methanoliparia (n=5)
c__Methanomicrobia (n=248)
c__Methanonatronarchaeia (n=6)
c__Methanopyri (n=5)
c__Methanosarcinia (n=254)
c__Poseidoniia (n=386)
c__SW−10−69−26 (n=5)
c__Syntropharchaeia (n=61)
c__Thermococci (n=116)
c__Thermoplasmata (n=553)
c__UBA148 (n=10)
c__UBA186 (n=3)
Euryarchaeota
c__Bathyarchaeia (n=285)
c__DRAE01 (n=1)
c__EX4484−205 (n=12)
c__JANJXX01 (n=10)
c__Korarchaeia (n=31)
c__Methanomethylicia (n=41)
c__Nitrososphaeria (n=505)
c__Nitrososphaeria_A (n=48)
c__QMWL01 (n=1)
c__Thermoprotei (n=89)
c__Thermoprotei_A (n=239)
TACK
0
25
50
75
100
no. of genomes (%)
Temperature
>80
60
45
30
15
0

**Figure S10: Estimated optimal growth temperature of archaea species representatives from the Genome Taxonomy Database r220.** Displayed is the proportion of genomes predicted to optimally grow within specific temperature ranges, using the GenomeSPOT v1.0.1 (Barnum et al. 2024). Rows on the x-axis represent archaeal classes grouped by their traditional supergroups: Asgardarchaeota, DPANN, Euryarchaeota and TACK. The x-axis indicates the percentage of tested archaeal reference genomes within each group.

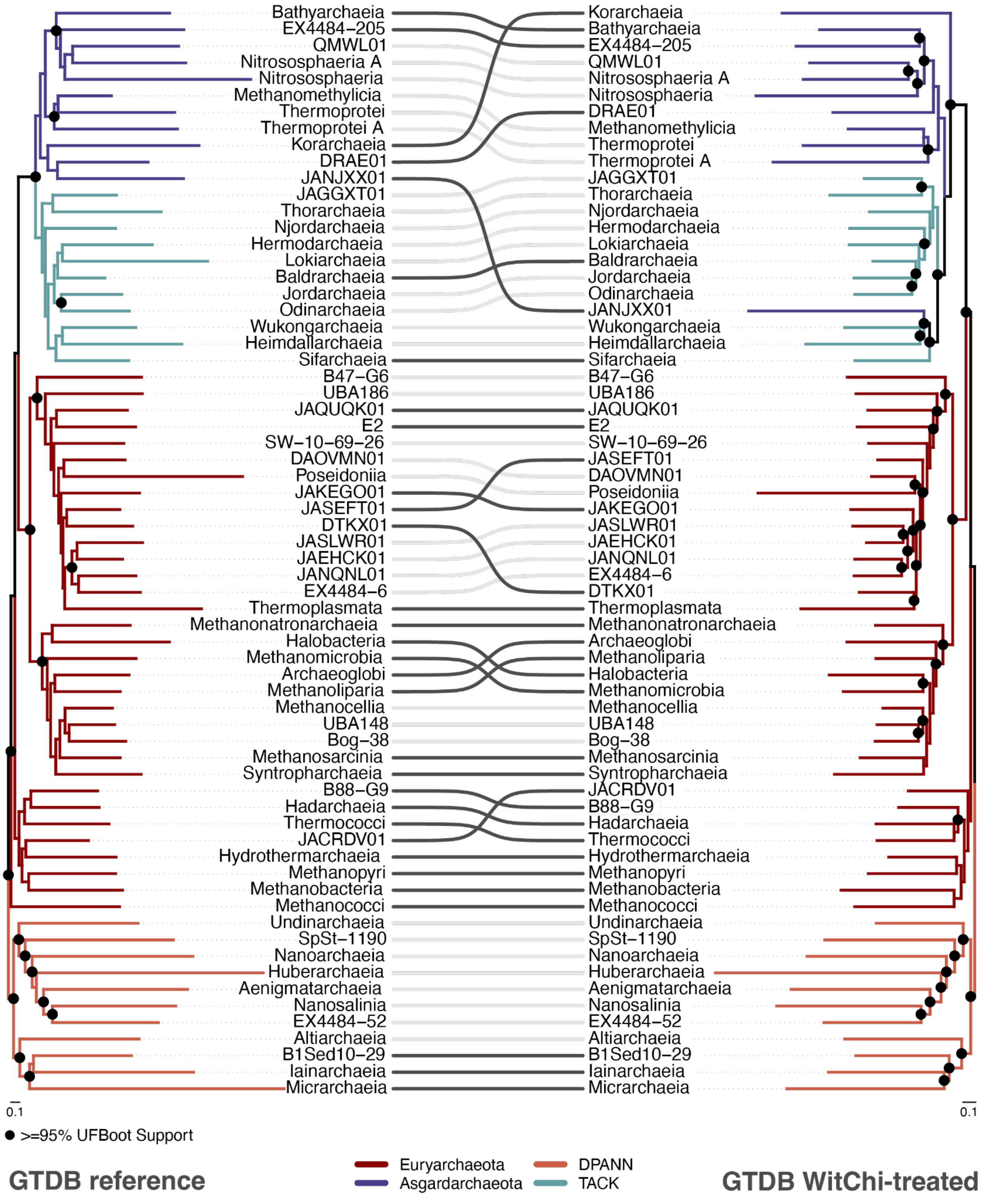


**Figure S11: Class-level tanglegram between the archaeal reference tree of Genome Taxonomy Database r220 (left) and with its WhitChi-treated counterpart (right).** The figure is based on the same dataset and data manipulation as Figure 4 of main text, but collapsed on class-level to display topological rearrangements within the different phyla. In the current release the Asgard class Atabeyarchaeia is misclassified as Njordarchaeia—an issue that has been corrected in the r226 release of the database. In short, the WitChi-treated phylogeny was inferred from the pruned alignment of 4,935 columns (down from 10,101). Light gray links indicate unchanged topology, while dark gray links highlight rearrangements. Both trees are rooted at the clade shown in orange, originally named after Diapherotrites, Parvarchaeota, Aenigmarchaeota, Nanoarchaeota and Nanohaloarchaeota (DPANN). Black circles at nodes indicate UFboot >= 95%. The scale bar represents the average number of substitutions per site.

**Table S1. Comparative overview of alignment pruning and compositional bias detection tools.**

| Software | Handles compositional heterogeneity | Removes phylogenetically uninformative sites | Reference |
|---|---|---|---|
| WitChi | $\chi^2$-based detection and pruning of alignment columns that drive taxon-specific composition bias using empirical null distributions that help for more accurate identification of biased taxa. | Indirectly. Biased columns are pruned, which may also remove low-signal sites. | Developed in this study |
| BMGE | Optional “stationary-based trimming” to remove columns causing compositional heterogeneity guided by pairwise comparisons of sequence composition. Also recoding options (RY coding for DNA, degenerate codons) to reduce bias. | Uses an entropy-like score per MSA column (weighted by similarity matrices like BLOSUM or PAM) to remove ambiguous or highly-variable columns. | (Criscuolo and Gribaldo 2010) |
| alignment_pruner.pl | $\chi^2$-based detection and pruning of alignment columns that drive taxon-specific composition bias but assuming independence among taxa (fixed $\chi^2$ threshold). Option to also remove taxa that are significantly deviating. | Removes gappy or unconserved columns using gap and conservation user-defined thresholds. | Used in: (Dharamshi et al. 2023; Huang et al. 2025; Martijn et al. 2018) |
| Phykit - compositional_bias_per_site | Diagnostic $\chi^2$ test per site quantifies compositional biased sites in an alignment. For each site, it performs a $\chi^2$ test on character distributions across taxa, returning the test statistic and corrected p-value to identify sites with significant compositional bias. | No but rather reports biased sites only and does not perform site removal. | (Steenwyk et al. 2024) |

| | | | |
|---|---|---|---|
| trimAl | Does not explicitly detect composition bias; trimming by similarity and gap structure may indirectly reduce bias. | Calculates per-column gap score and similarity score; trims based on user-defined thresholds or automatic modes (“gappyout”, “strict”, “strictplus”). | (Capella-Gutiérrez et al. 2009) |
| ClipKIT | Not designed for compositional bias but may indirectly reduce it by retaining informative sites. | Yes. Retains phylogenetically informative (e.g., parsimony-informative or constant) sites while discarding uninformative positions. | (Steenwyk et al. 2020) |
| Gblocks | Not designed for compositional bias detection; no explicit bias-aware trimming. | Selects conserved, contiguous alignment blocks based on minimum conservation and low gap density and discards poorly aligned and divergent regions. | (Castresana 2000) |

Criscuolo, Alexis, and Simonetta Gribaldo. 2010. “BMGE (Block Mapping and Gathering with Entropy): A New Software for Selection of Phylogenetic Informative Regions from Multiple Sequence Alignments.” *BMC Evolutionary Biology* 10 (July): 210.

Dharamshi, Jennah E., Stephan Köstlbacher, Max E. Schön, Astrid Collingro, Thijs J. G. Ettema, and Matthias Horn. 2023. “Gene Gain Facilitated Endosymbiotic Evolution of Chlamydiae.” *Nature Microbiology* 8 (1): 40–54.

Foster, Peter G., Dominik Schrempf, Gergely J. Szöllősi, Tom A. Williams, Cymon J. Cox, and T. Martin Embley. 2023. “Recoding Amino Acids to a Reduced Alphabet May Increase or Decrease Phylogenetic Accuracy.” *Systematic Biology* 72 (3): 723–737.

Huang, Wen-Cong, Maraike Probst, Zheng-Shuang Hua, et al. 2025. “Phylogenomic Analyses Reveal That Panguiarchaeum Is a Clade of Genome-Reduced Asgard Archaea Within the Njordarchaeia.” *Molecular Biology and Evolution* 42 (9). https://doi.org/10.1093/molbev/msaf201.

Martijn, Joran, Julian Vosseberg, Lionel Guy, Pierre Offre, and Thijs J. G. Ettema. 2018. “Deep Mitochondrial Origin Outside the Sampled Alphaproteobacteria.” *Nature* 557 (7703): 101–105.

Steenwyk, Jacob L., Thomas J. Buida 3rd, Yuanning Li, Xing-Xing Shen, and Antonis Rokas. 2020. “ClipKIT: A Multiple Sequence Alignment Trimming Software for Accurate Phylogenomic Inference.” *PLoS Biology* 18 (12): e3001007.

Steenwyk, Jacob L., Gemma I. Martínez-Redondo, Thomas J. Buida 3rd, et al. 2024. “PhyKIT: A Multitool for Phylogenomics.” *Current Protocols* 4 (11): e70016.
